# Reassessment of RNF43 Function Reveals No Impact on Endogenous EGFR or BRAF Protein Stability

**DOI:** 10.64898/2026.03.18.712374

**Authors:** Jiahui Niu, Shanshan Li, Ruyi Zhang, Jenna van Merode, Maikel P. Peppelenbosch, Ron Smits

**Author notes:** To whom correspondence should be addressed: Ron Smits, Erasmus MC, Dept. of Gastroenterology and Hepatology, room Na-1008, Wytemaweg 80, 3015 CN Rotterdam, The Netherlands.

## Abstract

RNF43 is best known for removing the Wnt-receptor complex from the cell surface, thereby maintaining Wnt-signaling at minimal essential levels. Recent studies reported that RNF43-mutant colorectal cancers carrying the common BRAFV600E mutation, respond more effectively to combined BRAF/EGFR inhibition. To determine whether RNF43 directly regulates EGFR or BRAF protein abundance, multiple pancreatic and colorectal cancer cell line models were generated in which RNF43 was knocked out, repaired, or stably overexpressed. Total and cell surface EGFR levels, as well as endogenous BRAF expression, were quantified.

Across all models, no consistent evidence emerges that RNF43 modulates endogenous EGFR or BRAF levels. R-spondins likewise fail to alter EGFR levels or internalization. Notably, elevated EGFR expression observed in a subset of RNF43 knockout clones is induced by unintended CRISPR/Cas9 vector integration rather than the absence of RNF43 itself, highlighting a previously underappreciated artefact that can confound interpretations of EGFR regulation in genome edited lines.

Overall, the data argue against a direct and general role for RNF43 in controlling EGFR or BRAF protein abundance, contradicting recent reports that propose degradation of these targets. Further studies are required to resolve these discrepancies and clarify the mechanistic basis underlying these conflicting observations.

## Introduction

Transmembrane E3 ubiquitin-protein ligases Ring Finger Protein 43 (RNF43) and Zinc And Ring Finger 3 (ZNRF3) are important regulators of the Wnt/β-catenin signaling pathway. They are frequently mutated in cancers, highlighting their crucial roles in tumorigenesis^[1–3]^. These proteins are best known for regulating the turnover of Wnt receptors, leading to their membrane clearance and subsequent downregulation of Wnt/β-catenin signaling. The membrane levels of RNF43/ZNRF3 themselves are, in turn, regulated by secreted roof plate-specific Spondins (R-spondin, RSPO) proteins^[1, 3, 4]^. Upon the formation of a complex between R-spondins, RNF43/ZNRF3, and LGR4/5/6 receptor proteins, RNF43 and ZNRF3 are cleared from the membrane. This facilitates the accumulation of Wnt receptors, thereby enhancing cellular responsiveness to Wnt-ligand activation.

More recently, RNF43/ZNRF3 have also been identified as important regulators in a variety of other signaling pathways, indicating their broader influence beyond just the traditional Wnt/β-catenin pathway^[3, 5]^. Examples include non-canonical WNT/PCP signaling^[6, 7]^, and the inhibition of PAR2-enhanced β-catenin signaling by promoting the ubiquitination and degradation of membrane-bound PAR2^[8]^. Furthermore, research from Christof Niehrs’ group showed that RNF43/ZNRF3 can induce the endocytosis and degradation of bone morphogenetic protein receptor type 1A (BMPR1A), one of the receptors responsible for transducing BMP signals^[9–11]^. This downregulation requires the simultaneous binding of RSPO2 or RSPO3 to both extracellular BMPR1A and RNF43/ZNRF3. Additionally, RSPO2 has been found to promote the internalization of fibroblast growth factor receptor 4 (FGFR4) via ZNRF3-mediated endocytosis^[12]^. All these findings reveal that RNF43/ZNRF3, sometimes in conjunction with R-spondins, can influence not only Wnt/β-catenin signaling but also various other signaling pathways originating from the cell membrane.

Colorectal cancer (CRC) is the third most common cancer and the second leading cause of cancer-related deaths globally^[13]^. Although survival rates have improved, metastatic CRC (mCRC) remains highly lethal, with a 5-year survival rate of approximately 14%^[14]^. Most mCRCs are microsatellite stable (MSS) and typically do not respond well to checkpoint inhibitors, resulting in less favorable treatment outcomes. There is still some uncertainty about what the best first-line therapy should be for these patients^[15]^. BRAF is a member of the RAF protein family involved in the EGFR-mediated MAPK pathway. Approximately 10% of advanced mCRC cases acquire the BRAF^V600E^ mutation, which is associated with a particularly aggressive disease phenotype^[16]^. Despite their effectiveness in melanoma, BRAF inhibitors alone or in combination with MEK inhibitors have shown limited clinical benefit in patients with mCRC^BRAF-V600E^, likely due to adaptive feedback activation of the EGFR-mediated MAPK signaling pathway^[17–19]^.

In 2022, a large-scale genetic biomarker analysis was conducted on mCRC^BRAF-V600E^ patients treated with anti-BRAF/EGFR combination therapy. This analysis identified *RNF43* mutations in 29% of the MSS-mCRC^BRAF-V600E^ cases within their dataset, and revealed that these inactivating *RNF43* mutations were linked to improved response rates and better survival outcomes in patients with MSS-mCRC^BRAF-V600E^ ^[20]^, a finding that was later confirmed in two independent studies^[21, 22]^. The underlying mechanism for this beneficial effect of *RNF43* mutation on patient outcome was however not addressed.

Based on this study and the involvement of RNF43/ZNRF3 in regulating other (membrane) components besides the Wnt-receptor, we hypothesized that RNF43 might directly influence the activity of EGFR and/or BRAF. While our research was ongoing, Yue and colleagues published a preprint on bioRxiv indeed suggesting that RNF43/ZNRF3 are novel E3 ubiquitin ligases for EGFR, and established the inactivation of ZNRF3/RNF43 as a key factor driving enhanced EGFR signaling. RNF43/ZNRF3 were shown to bind to EGFR through their extracellular R-spondin/PA domains, and subsequent ubiquitination and degradation required the RING domain^[23]^. Additionally, another recent study demonstrated that RNF43 targets BRAF as a substrate in vivo, resulting in ubiquitination at position K499 and subsequent BRAF degradation via the proteasome^[24]^. This indicates that RNF43 may play a role in suppressing tumor growth by inhibiting the BRAF/MEK signaling pathway. If both these findings are correct, it provides a potential rationale for the increased sensitivity of RNF43/BRAF mutant colorectal cancers to combined EGFR/BRAF inhibition. RNF43 mutation not only enhances Wnt-induced β-catenin signaling but would also elevate levels of EGFR and mutant BRAF. Consequently, these cancers may become more addicted to increased EGFR/BRAF signaling for their growth, rendering them more responsive to EGFR/BRAF inhibition.

Here, we conducted an independent investigation into the proposed role of RNF43 in regulating EGFR and BRAF protein levels. Our findings do not support the claims made in these recent publications, casting doubt on whether these explanations are valid for the increased sensitivity of BRAF/RNF43-mutant colorectal cancers to combined BRAF/EGFR inhibition.

## Results

### 3.1 Baseline characteristics of employed cell lines and RNF43-altered clones thereof

To investigate the potential interactions between RNF43 and protein levels of EGFR and BRAF, we analyzed three cancer cell lines: AsPC-1 (pancreatic ductal adenocarcinoma), Caco-2 (colon adenocarcinoma), and HT-29 (colon adenocarcinoma). Table 1 summarizes the key mutations observed in these cell lines for β-catenin related genes, BRAF, and their microsatellite stability status. Notably, AsPC-1 harbors a homozygous p.S720* truncating mutation in RNF43 and a previously unrecognized heterozygous p.E62* mutation in ZNRF3 (Supplementary Figure S1). Both Caco-2 and HT-29 carry inactivating APC mutations, with HT-29 additionally harboring the common BRAF^V600E^ mutation. All three cell lines possess a functional mismatch repair system.

**Table 1.** Cell lines used in this study. Relevant mutations and microsatellite stability (MSS) status are depicted. homo., homozygous; het., heterozygous; wt., wildtype, KO, Knockout; OE, Overexpression.

| Cells | Disease | Relevant mutations in parental cells |  |  |  |  |  | RNF43 altered clones |  |  |
| --- | --- | --- | --- | --- | --- | --- | --- | --- | --- | --- |
|  |  | RNF43 | ZNRF3 | APC | CTNNB1 | BRAF | MSI status | KO | OE | Repaired |
| AsPC-1 | Pancreatic ductal adenocarcinoma | p.S720* homo. | p.E62* het. | wt | wt | wt | MSS | A1<br>A5<br>C1 | - | 5E<br>5F<br>5G |
| Caco-2 | Colon adenocarcinoma | wt | wt | p.Q1367* homo. | p.G245A het. | wt | MSS | D1<br>F3<br>F5 | A10<br>C3 | - |
| HT-29 | Colon adenocarcinoma | wt | wt | p.E853* het.+<br>p.T1556fs*3 het. | wt | p.V600E het.+<br>p.T119S het. | MSS | A3<br>B1 | - | - |

To further investigate the functional consequences of RNF43 alterations, we initially generated 2-3 independent RNF43-knockout (KO) clones for each cell line (Supplementary Figure S2). We also established two Caco-2 clones stably overexpressing FLAG-tagged RNF43 protein, with RNA expression levels about 8-fold and 25-fold higher than endogenous *RNF43* (Supplementary Figure S3). As the AsPC-1 cell line harbors a homozygous S720* RNF43 mutation that may have a partial loss-of-function, we also generated isogenic clones in which this mutation was corrected to wild-type *RNF43* (Supplementary Figure S4). Functional assays were performed in parental, RNF43-repaired, and KO AsPC-1 clones using a β-catenin luciferase reporter assay. As expected, β-catenin signaling was highest in KO clones, reduced in parental cells, and further suppressed in repaired clones (Supplementary Figure S5). The successfully derived clones are listed in Table 1, with multiple independent clones established for each condition.

### 3.2 Analysis of EGFR protein levels in relation to RNF43 functionality

EGFR expression and activation, as well as downstream signaling, were first assessed in the AsPC-1 cell line across parental cells, RNF43-repaired clones, and RNF43-knockout clones under both basal conditions and EGF stimulation. RNF43 status did not appear to significantly influence total EGFR levels (Figure 1A). Similarly, a short-term 30 minutes EGF treatment did not noticeably alter total EGFR levels (Figure 1A). Given that short-term EGF stimulation might not reveal long-term regulatory effects, EGF treatment was extended to 48 hours (Supplementary Figure S6). Although total EGFR levels were markedly reduced following extended EGF stimulation, again no clear correlation with RNF43 functionality was detected.

**Fig 1.**
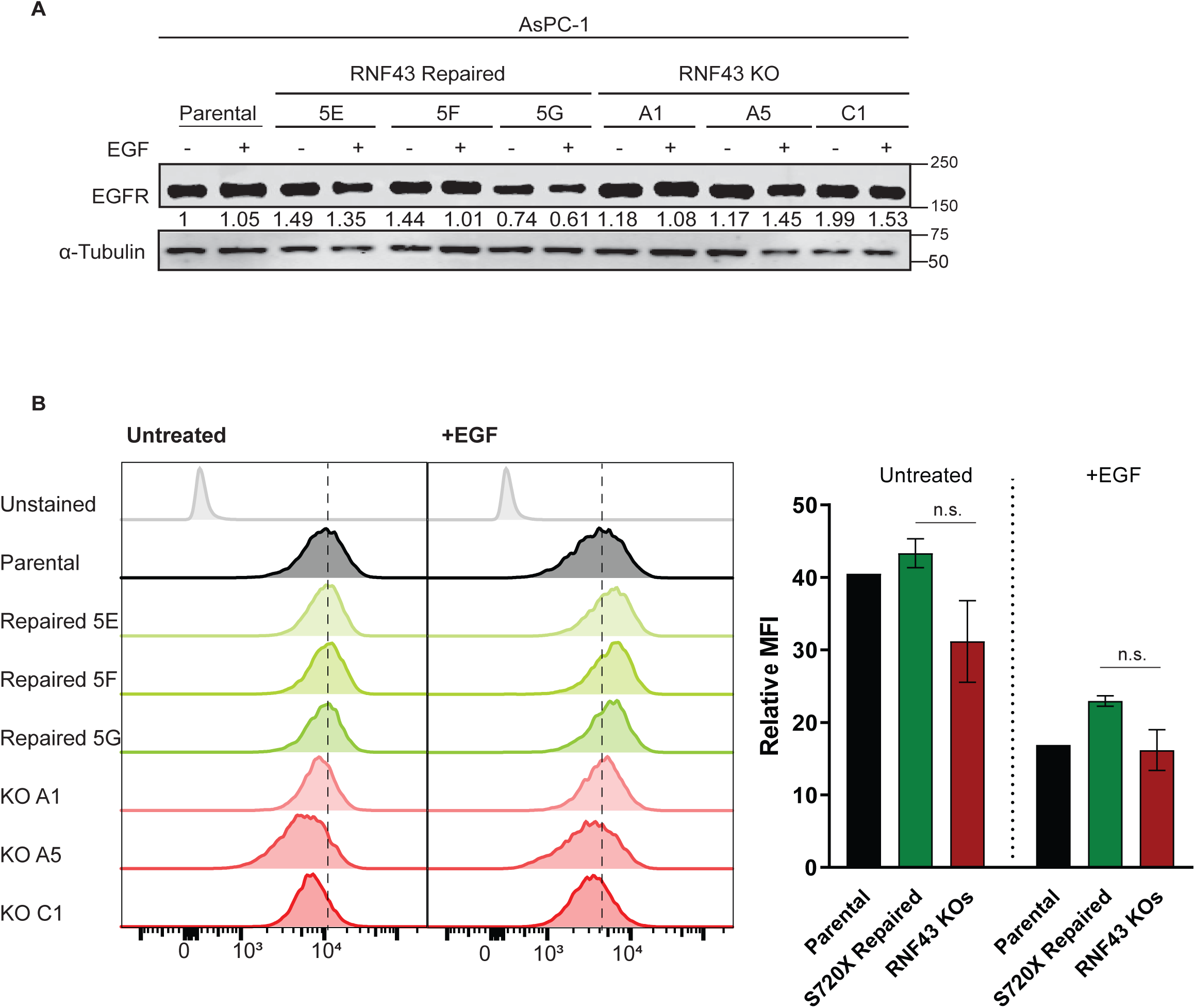
RNF43 status does not correlate with total or cell-surface EGFR levels in AsPC-1 cells. (A) Immunoblot analysis of total EGFR in parental AsPC-1 cells, RNF43-repaired clones (5E, 5F, 5G), and RNF43 knockout clones (A1, A5, C1) under basal conditions (−EGF) and after 30 min EGF stimulation (+EGF). α-Tubulin served as a loading control. Numbers below the EGFR bands indicate relative EGFR levels normalized to α-tubulin and expressed relative to the parental −EGF condition (set to 1). (B) Flow cytometry analysis of cell-surface EGFR in parental AsPC-1 cells and RNF43 altered clones under basal conditions (−EGF) or after 30 min EGF stimulation (+EGF). Representative histogram overlays are shown (left). The bar graph (right) summarizes mean fluorescence intensity (MFI) for parental cells and for RNF43-repaired clones (n = 3) and RNF43 KO clones (n = 3). Bars show mean ± SD across clones. Data shown are representative of three independent experiments. Statistical significance was assessed separately for the −EGF and +EGF conditions using Welch’s one-way ANOVA, followed by Dunnett’s T3 multiple-comparisons test comparing each group to parental; n.s., not significant.

Although this analysis suggests that RNF43 has no effect on total EGFR levels in AsPC-1 cells, it might still affect cell-surface levels of EGFR. To explore this possibility, we therefore quantified surface EGFR by flow cytometry. As shown in Figure 1B, EGFR membrane levels remained unchanged across all RNF43-repaired clones, while a slight decrease was observed in knock-out clones. This latter finding contradicts expectations if RNF43 is capable of reducing EGFR levels. As expected, short-term EGF treatment uniformly reduced EGFR membrane levels across all clones.

A similar analysis was performed in all Caco-2 and HT-29 clones. No alteration in total or membrane EGFR levels is observed in both Caco-2 overexpression clones (Figure 2), while a 1.6-1.75 fold increase is observed in two out of three knockout clones. Among the HT-29 knockout clones, clone B1 exhibited a 1.6-fold increase in EGFR levels (Figure 3), whereas clone A3 did not show this trend. Supplementary Figure S7 presents changes in phosphorylated EGFR and downstream pERK1/2 following EGF addition across all clones, which largely align with expectations.

**Fig 2.**
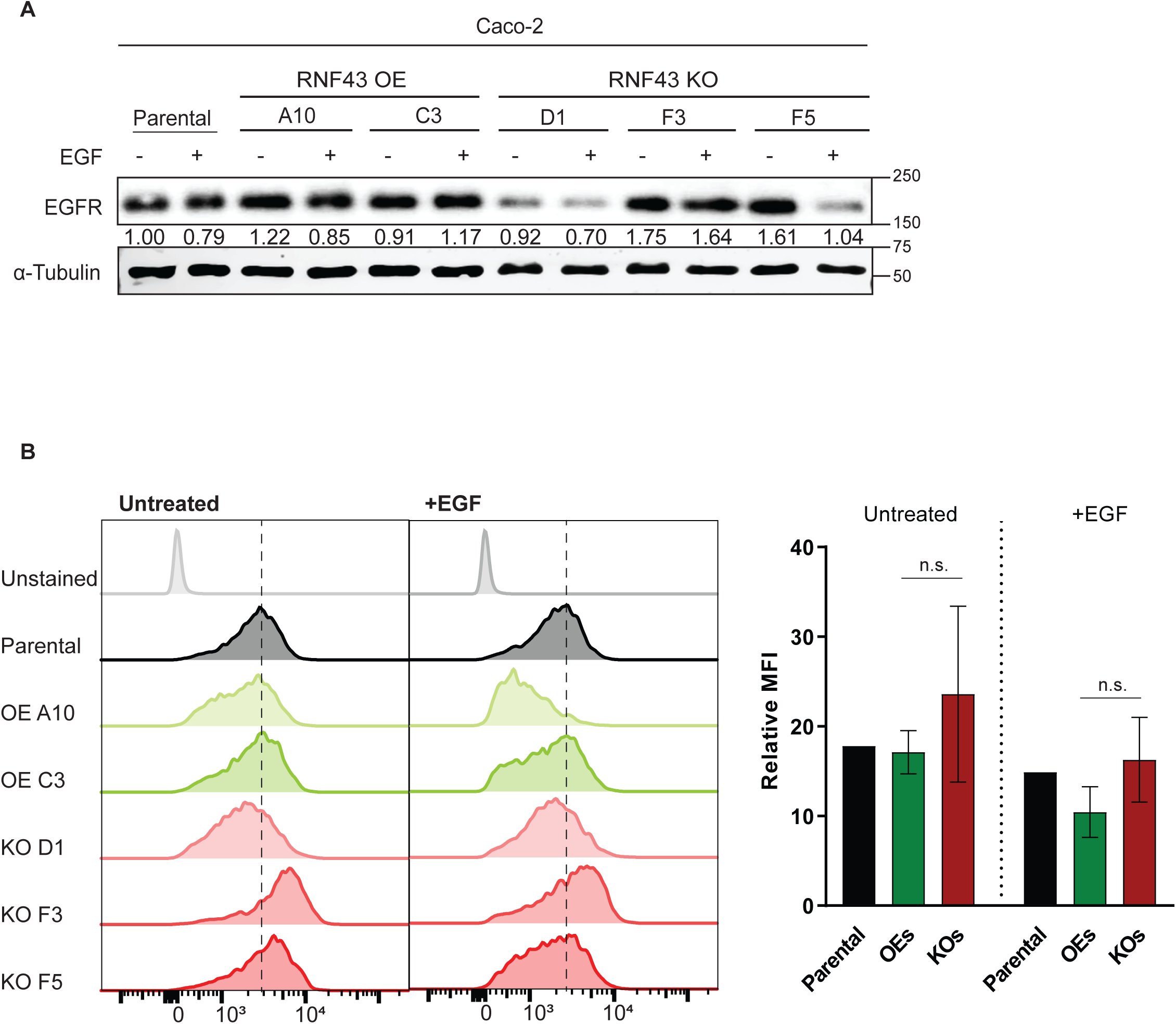
RNF43 status does not correlate with total or cell-surface EGFR levels in Caco-2 cells. (A) Immunoblot analysis of total EGFR in parental Caco-2 cells, RNF43-overexpressing clones (A10, C3), and RNF43 knockout clones (D1, F3, F5) under basal conditions (−EGF) and after 30 min EGF stimulation (+EGF). α-Tubulin served as a loading control. Numbers below the EGFR bands indicate relative EGFR levels normalized to α-tubulin and expressed relative to the parental −EGF condition (set to 1). (B) Flow cytometry analysis of cell-surface EGFR in parental Caco-2 cells and RNF43 altered clones under basal conditions (−EGF) or after 30 min EGF stimulation (+EGF). Representative histogram overlays are shown (left). The bar graph (right) summarizes mean fluorescence intensity (MFI) for parental cells and for RNF43 OE clones (n = 2) and RNF43 KO clones (n = 3). Bars show mean ± SD across clones. Data shown are representative of three independent experiments. Statistical significance was assessed by Welch’s one-way ANOVA performed separately for the −EGF and +EGF conditions, followed by Dunnett’s T3 multiple-comparisons test comparing each group to parental; n.s., not significant.

**Fig 3.**
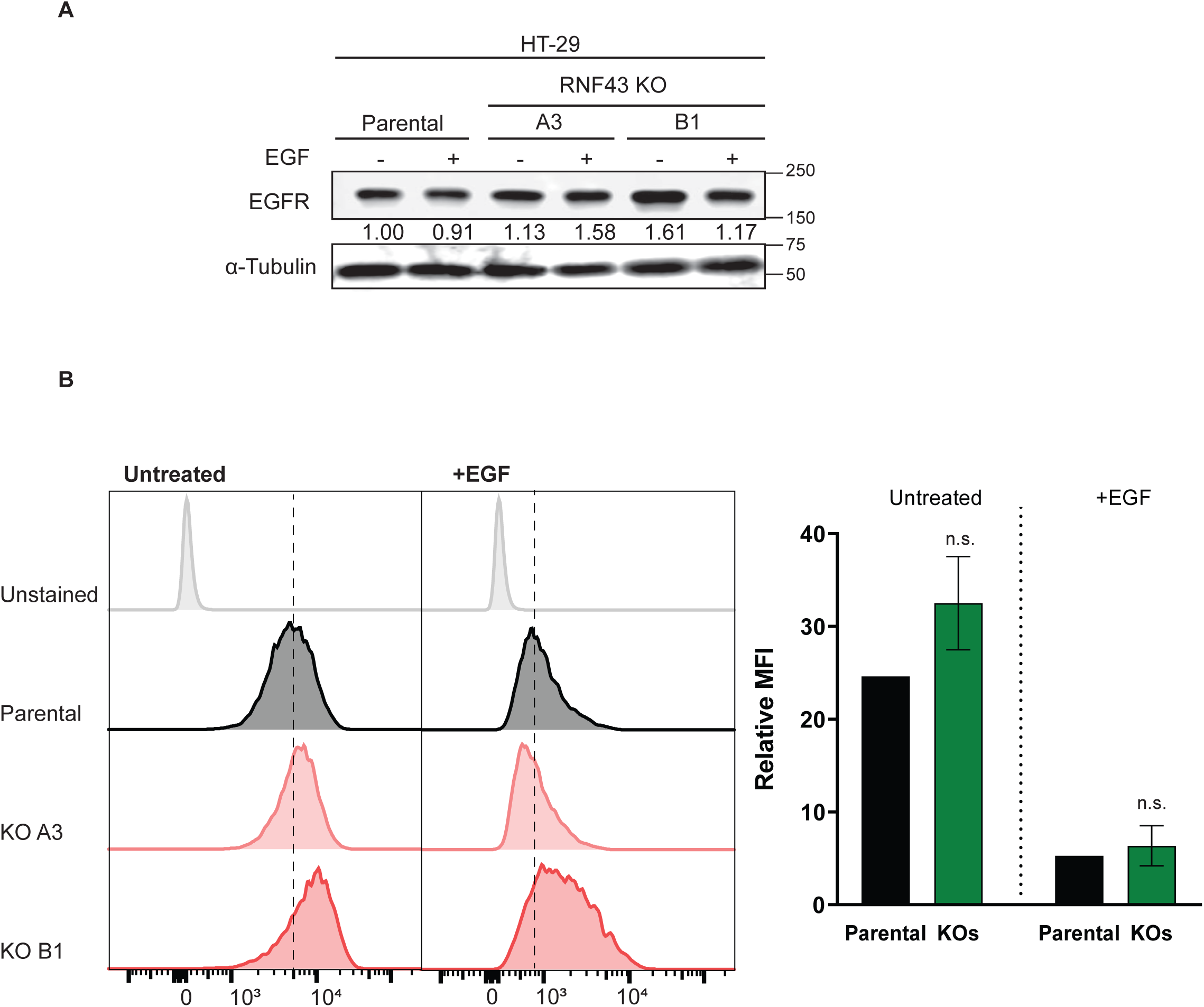
RNF43 knockout does not consistently alter total or cell-surface EGFR levels in HT-29 cells. (A) Immunoblot analysis of total EGFR in parental HT-29 cells and RNF43 knockout clones (A3, B1) under basal conditions (−EGF) and after 30 min EGF stimulation (+EGF). α-Tubulin is shown as a loading control. Numbers below the EGFR bands indicate relative EGFR levels normalized to α-tubulin and expressed relative to the parental −EGF condition (set to 1). (B) Flow cytometry analysis of cell-surface EGFR in parental HT-29 cells and RNF43 KO clones under basal conditions (−EGF) or after 30 min EGF stimulation (+EGF). Representative histogram overlays are shown (left). The bar graph (right) summarizes mean fluorescence intensity (MFI) for parental cells and for RNF43 KO clones (n = 2); bars show mean ± SD across clones. Data shown are representative of three independent experiments. Statistical significance was assessed separately for the −EGF and +EGF conditions using Welch’s unpaired two-tailed t-test; n.s., not significant.

Overall, we did not observe a correlation between RNF43 functionality and EGFR levels in AsPC-1 and RNF43-overexpressing Caco-2 clones, while a moderate increase is observed in some Caco-2 and HT-29 knockout clones. These latter clones were generated using a short-term puromycin selection to avoid stable integration of the pX459 Cas9 genome editing vector. Nevertheless, two Caco-2 clones showing moderate EGFR upregulation had inadvertently integrated this construct, as observed by expression of FLAG-tagged Cas9 (Supplementary Figure S8).

We hypothesized that sustained expression of Cas9 and/or other components of the pX459 vector might influence cellular processes, including EGFR regulation. To investigate this, we generated multiple independent HT-29 clones, varying in pX459 integration status and RNF43 knockout. Immunoblot analysis unexpectedly showed that all Cas9-expressing clones exhibited increased EGFR levels, an effect not observed in the RNF43 knockout clones without pX459 integration (Figure 4A). These findings indicate that the pX459 vector itself may aberrantly elevate EGFR expression. Consistent with this, we generated additional Caco-2 clones with stable pX459 integration and sustained Cas9 expression and likewise observed increased EGFR levels relative to Cas9-negative controls (Figure 4B).

**Fig 4.**
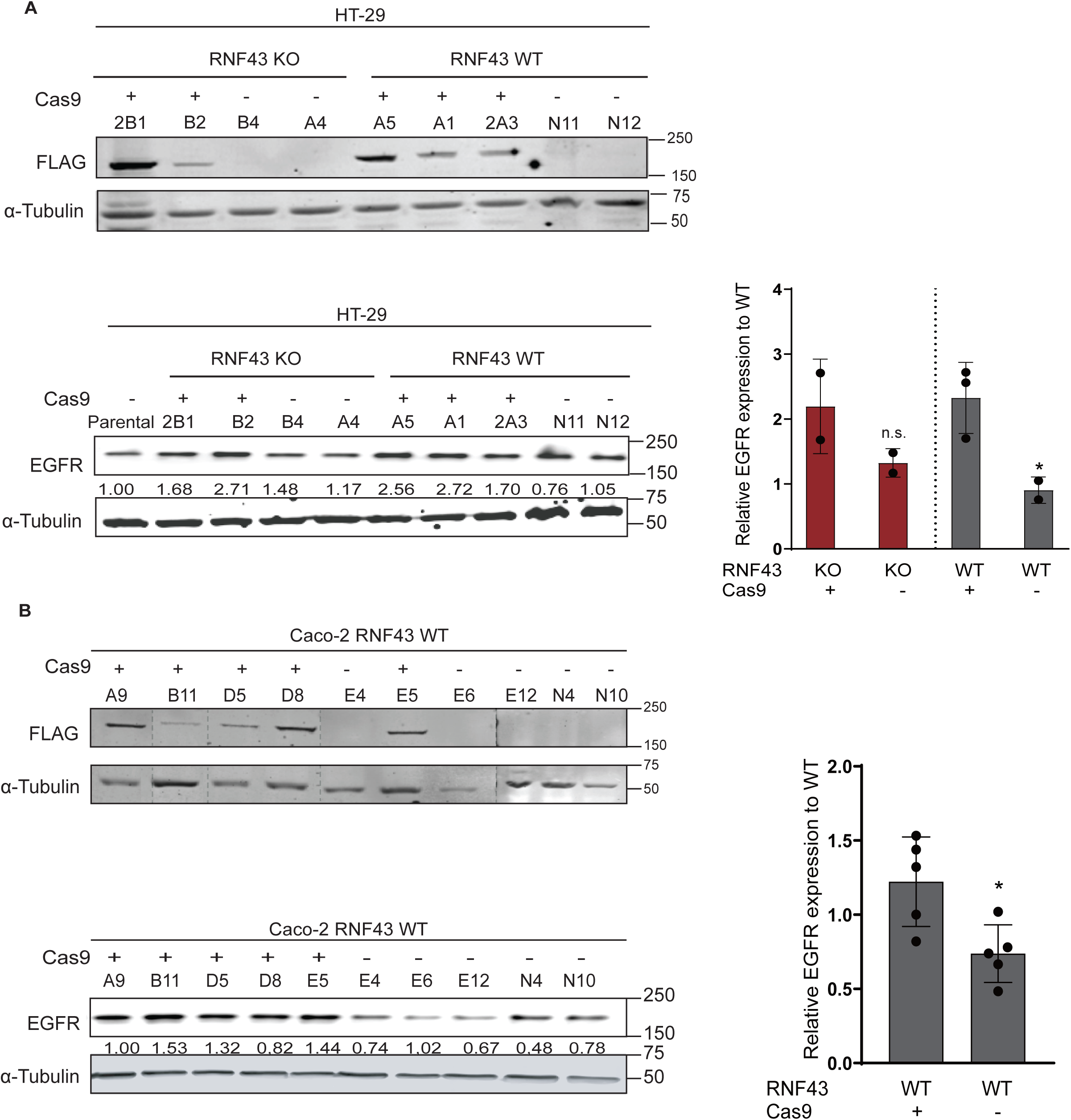
Sustained Cas9 expression from pX459 is associated with increased EGFR levels in HT-29 and Caco-2 clones, independent of RNF43 status. (A) HT-29 clones differing in RNF43 genotype (RNF43 KO vs RNF43 WT) and Cas9 status were generated by transfection with pX459 carrying an RNF43-targeting gRNA (KO) or empty pX459 (WT controls), followed by puromycin selection and single-colony isolation, as described in Materials and Methods. The upper immunoblot shows FLAG-tagged Cas9 to stratify clones as Cas9-positive or Cas9-negative; α-tubulin served as a loading control. The lower immunoblot shows total EGFR levels in the same HT-29 clone panel. Numbers below the EGFR bands indicate EGFR band intensities normalized to α-tubulin and expressed relative to the parental sample (set to 1). The bar graph summarizes relative EGFR expression grouped by RNF43 genotype and Cas9 status; dots represent individual clones and bars show mean ± SD across clones. Statistical significance was assessed by Welch’s unpaired two-tailed t-test comparing Cas9-positive versus Cas9-negative clones within each RNF43 genotype; n.s., not significant; *P < 0.05. (B) Caco-2 RNF43 WT clones were generated by transfection with empty pX459 followed by puromycin selection and single-colony isolation, as described in Materials and Methods. The upper immunoblot shows FLAG-tagged Cas9 to stratify clones as Cas9-positive or Cas9-negative; α-tubulin served as a loading control. The lower immunoblot shows total EGFR levels in the same Caco-2 clone panel. Numbers below the EGFR bands indicate EGFR band intensities normalized to α-tubulin and expressed relative to clone A9 (set to 1). The bar graph summarizes relative EGFR expression in Cas9-negative versus Cas9-positive clones; dots represent individual clones and bars show mean ± SD across clones. Data shown are representative of three independent experiments. Statistical significance was assessed by Welch’s unpaired two-tailed t-test; *P < 0.05. Where indicated by dashed lines, lanes were juxtaposed from non-adjacent positions on the same blot for presentation.

### 3.3 Transient overexpression of RNF43 or ZNRF3 does not alter EGFR levels

Thus far, our results do not show a clear link between RNF43 functionality and EGFR total or membrane levels. This contrasts with a recent bioRxiv preprint, which proposes that loss of ZNRF3/RNF43 leads to increased EGFR levels and activity in cancer ^[23]^. Therefore, we decided to repeat one of their more straightforward experiments, that is to transiently co-express EGFR with either RNF43 or ZNRF3 in HEK293T cells. We also included RNF43/ZNRF3 variants with RING domain mutations that disrupt their ubiquitination functionality^[25]^, serving as negative controls. No significant differences in EGFR expression were observed among the different constructs, suggesting that neither RNF43 nor ZNRF3 directly regulate EGFR protein levels under these experimental conditions (Figure 5A). However, in line with expectations, both wild-type RNF43 and ZNRF3 effectively reduced mature FZD5 levels (Figure 5B)^[26–29]^.

**Fig 5.**
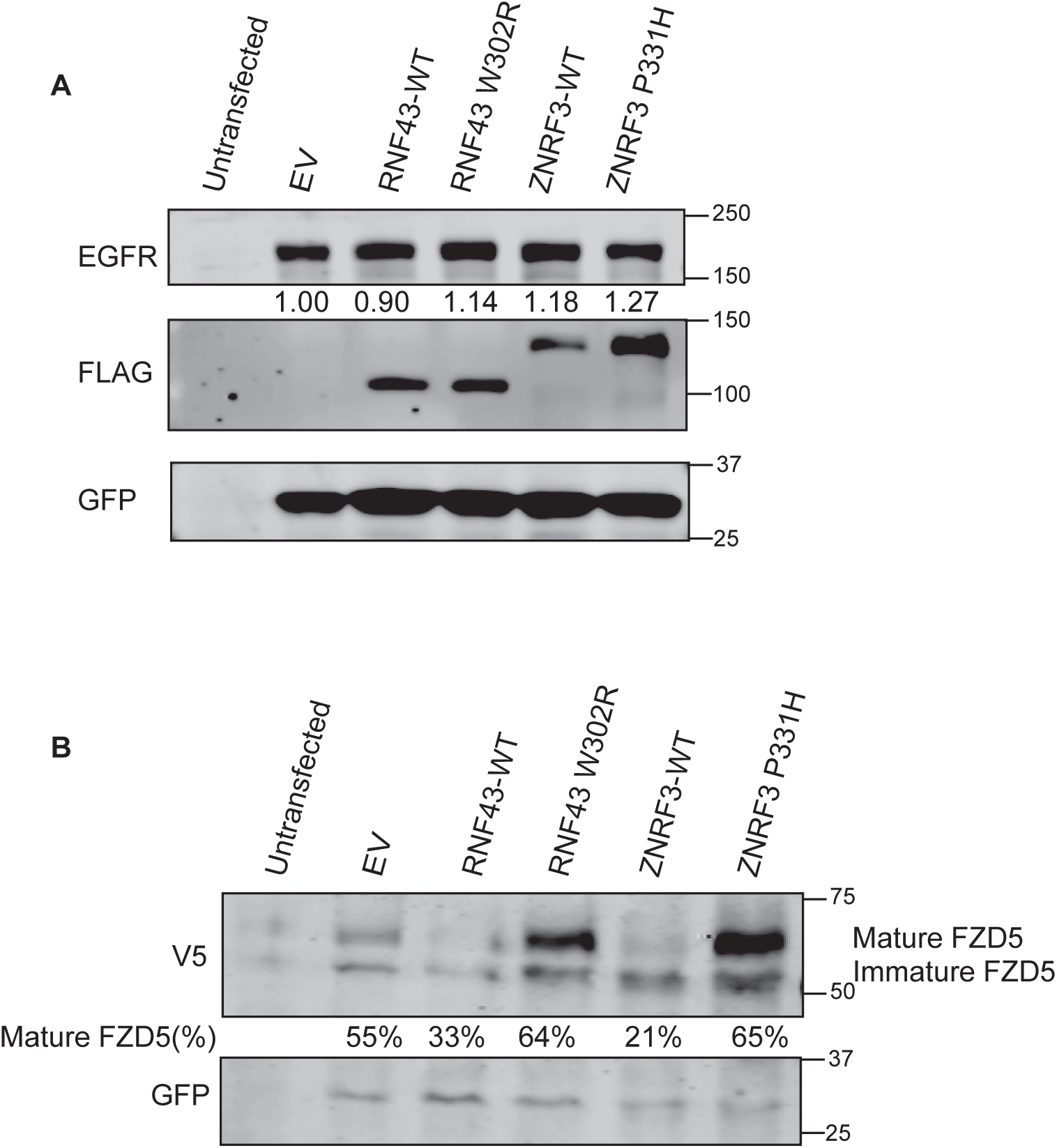
RNF43/ZNRF3 overexpression does not markedly alter total EGFR levels, with FZD5 processing used as a functional control. (A) Immunoblot analysis of total EGFR in HEK293T cells left untransfected or transfected with empty vector (EV), FLAG-tagged RNF43-WT, RNF43 W302R, ZNRF3-WT, or ZNRF3 P331H. FLAG immunoblotting confirms expression of the indicated RNF43/ZNRF3 constructs, and GFP is shown as a transfection control. Numbers below the EGFR bands indicate relative EGFR levels normalized to GFP and expressed relative to EV (set to 1). (B) Immunoblot of V5-tagged FZD5 in HEK293T cells, included as a functional readout to confirm RNF43/ZNRF3 activity under the same transfection conditions as in (A). Mature and immature FZD5 species are indicated, and the percentage of mature FZD5 (relative to total FZD5 signal) is shown below each lane. GFP is shown as a transfection control. Data shown are representative of three independent experiments.

### 3.4 No evidence of EGFR internalization after RSPO treatment

Previous studies identified RSPO2 and RSPO3 as high-affinity ligands for the cell-surface receptor BMPR1A, capable of forming a ternary complex with RNF43/ZNRF3 to regulate BMP receptor endocytosis and degradation^[10, 11]^. Based on this, we hypothesized that a similar mechanism might apply to the EGF receptor, and therefore we introduced RSPO proteins in our investigation. For this purpose, we generated L-cell clones that stably overexpress RSPO1, RSPO2, RSPO3, and two isoforms of RSPO4 (Supplementary Figure S9), and additionally utilized a HEK293T clone expressing mouse RSPO1 commonly used in organoid cultures.

EGFR flow cytometry performed on parental Caco-2 and HT-29 cells revealed no substantial changes in EGFR membrane levels after 48 hours of exposure to these different RSPO-conditioned media (Figure 6A, B). In contrast, prolonged EGF treatment resulted in a near-complete depletion of membrane EGFR. Similar results were obtained in the RNF43-overexpressing Caco-2 clone C3, which showed no detectable alterations in EGFR levels upon RSPO-treatment (Figure 6C).

**Fig 6.**
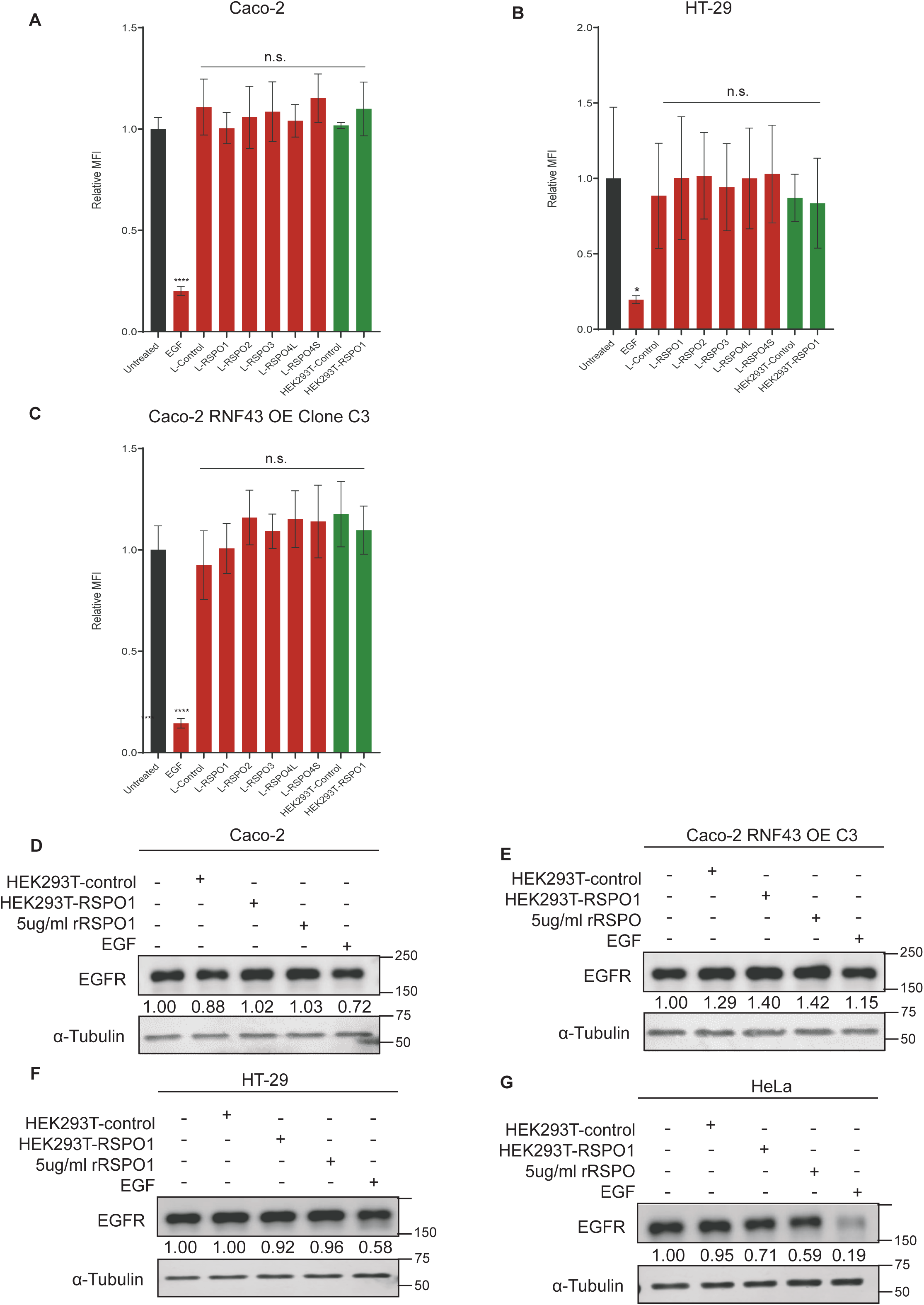
RSPO stimulation does not increase total or cell-surface EGFR levels across multiple cell lines. (A–C) Flow cytometry analysis of cell-surface EGFR in Caco-2 (A), HT-29 (B), and Caco-2 RNF43 OE clone C3 (C) cells following 48 h treatment with the indicated RSPO-conditioned media or controls. rRSPO1 was used at 5 µg/ml. All conditioned media were used at a 1:10 dilution. Cell-surface EGFR is shown as relative mean fluorescence intensity (MFI) normalized to the untreated condition (set to 1). Bars show mean ± SD across replicate measurements. EGF treatment was included as a positive control for EGFR downregulation. (D–G) Immunoblot analysis of total EGFR in Caco-2 (D), Caco-2 RNF43 OE clone C3 (E), HT-29 (F), and HeLa (G) cells treated for 3 h with HEK293T-control conditioned medium, HEK293T-RSPO1 conditioned medium, rRSPO1 (5 µg/mL), or EGF, as indicated. α-Tubulin served as a loading control. Numbers below the EGFR bands indicate relative EGFR levels normalized to α-tubulin and expressed relative to the untreated condition (set to 1). Data shown are representative of three independent experiments. Statistical significance was assessed using Welch’s one-way ANOVA, followed by Dunnett’s T3 multiple-comparisons test comparing each group to the untreated control; *P < 0.05, ****P < 0.0001, n.s., not significant.

Importantly, this experiment also indirectly supports our observation that RNF43 and ZNRF3 do not play a significant role in reducing membrane EGFR levels. R-spondins effectively remove RNF43 and ZNRF3 from the membrane, essentially mimicking a functional knockout of both proteins, a condition expected to increase EGFR levels at the membrane. Since no such increase was observed, our findings further reinforce that RNF43 and ZNRF3 do not regulate EGFR levels.

Similar experiments were performed in the recent bioRxiv preprint by Yue et al.^[23]^. A 2-4 hour recombinant RSPO2 treatment enhanced EGFR protein levels in HT-29 cells, while RSPO1 increased EGFR expression in *Apc*^min^ mouse intestinal tumor organoids. To assess whether a similar effect occurs in our case, we analyzed total EGFR expression in Caco-2 cells, an RNF43-overexpressing clone, as well as HT-29 and HeLa cells following a 3-hour treatment with either RSPO1 derived from HEK293T-conditioned medium or recombinant RSPO2 protein. Under all tested conditions, total EGFR levels remained unchanged compared to controls (Figure 6D–G).

To complement these findings with subcellular resolution, we performed EGFR immunofluorescence microscopy on parental Caco-2 and an RNF43-overexpressing clone, as well as parental AsPC-1 cells alongside RNF43-repaired and knockout AsPC-1 clones. Cells were treated with EGF (50 ng/mL) or individual RSPO isoforms (RSPO1–4L) for 30 minutes. EGF exposure led to a robust redistribution of EGFR from the plasma membrane to intracellular vesicles in all lines (Figure 7, Supplementary Figure S10), while RSPO-treated cells retained mostly membrane-localized EGFR with some intracellular vesicles, indistinguishable from untreated controls. Importantly, this pattern held across all RNF43 genotypes, that is wild-type, overexpressed, repaired, and knockout, demonstrating that neither RSPO isoforms nor RNF43 perturbation alters EGFR subcellular localization patterns.

**Fig 7.**
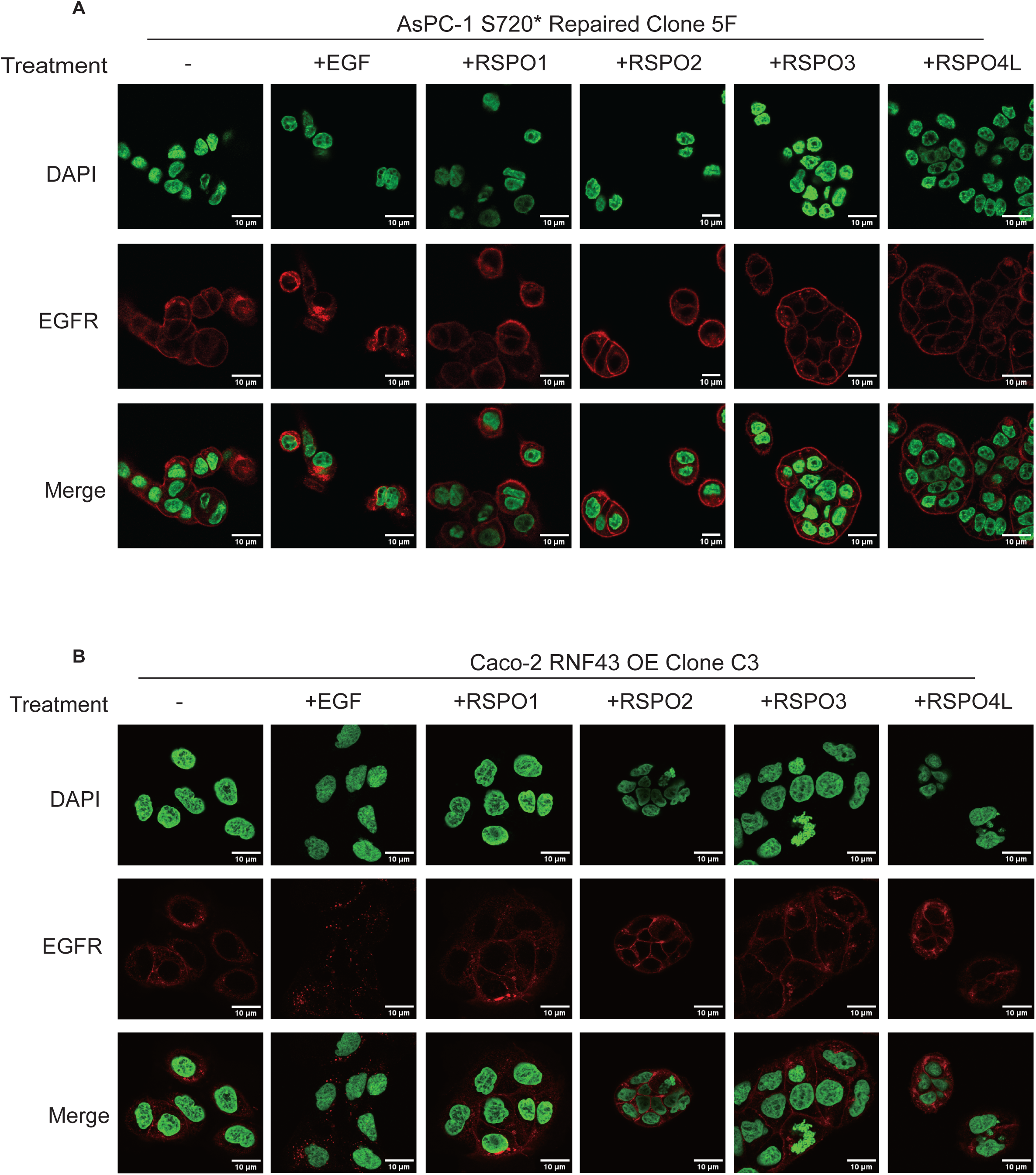
RSPO stimulation does not affect internalization of EGFR in Caco-2 or AsPC-1 cells. Immunofluorescence staining of EGFR following 30 min treatment with EGF (+EGF) or RSPO-conditioned media containing RSPO1, RSPO2, RSPO3, or RSPO4L (+RSPO1/2/3/4L), as indicated. Images are shown for AsPC-1 S720* repaired clone 5F (A), Caco-2 RNF43 OE clone C3 (B). EGFR is shown in red and nuclei (DAPI) in green; merged images are shown in the bottom row for each condition. Scale bars, 10 μm.

### 3.5 RNF43 functionality does not appear to correlate with BRAF protein levels

Thus far, our results do not support our hypothesis that RNF43 plays a role in regulating EGFR functionality, which could have provided an explanation why *RNF43* mutation may improve the response of BRAF-mutant colorectal cancers to combined BRAF/EGFR inhibition^[20–22]^. A second alternative hypothesis is that RNF43 influences BRAF protein levels. Support for this hypothesis comes from a recent report stating that RNF43 directly ubiquitinates both wild-type and V600E-mutant BRAF, leading to their degradation via the proteasome^[24]^. We used our cell clone collection to find further support for this observation. However, in none of the RNF43 knockout clones did we observe the expected increase in BRAF levels (Figure 8). Likewise, we did not observe a reduction in BRAF levels in either the RNF43-repaired AsPC-1 or the RNF43-overexpressing Caco-2 clones. Using HEK293T cells, we also co-expressed FLAG-tagged wild-type or V600E-mutant BRAF with either HA-tagged RNF43 or ZNRF3 (Supplementary Figure S11A). For both BRAF variants, we observed at most a slight reduction when co-expressed with RNF43 or ZNRF3.

**Fig 8.**
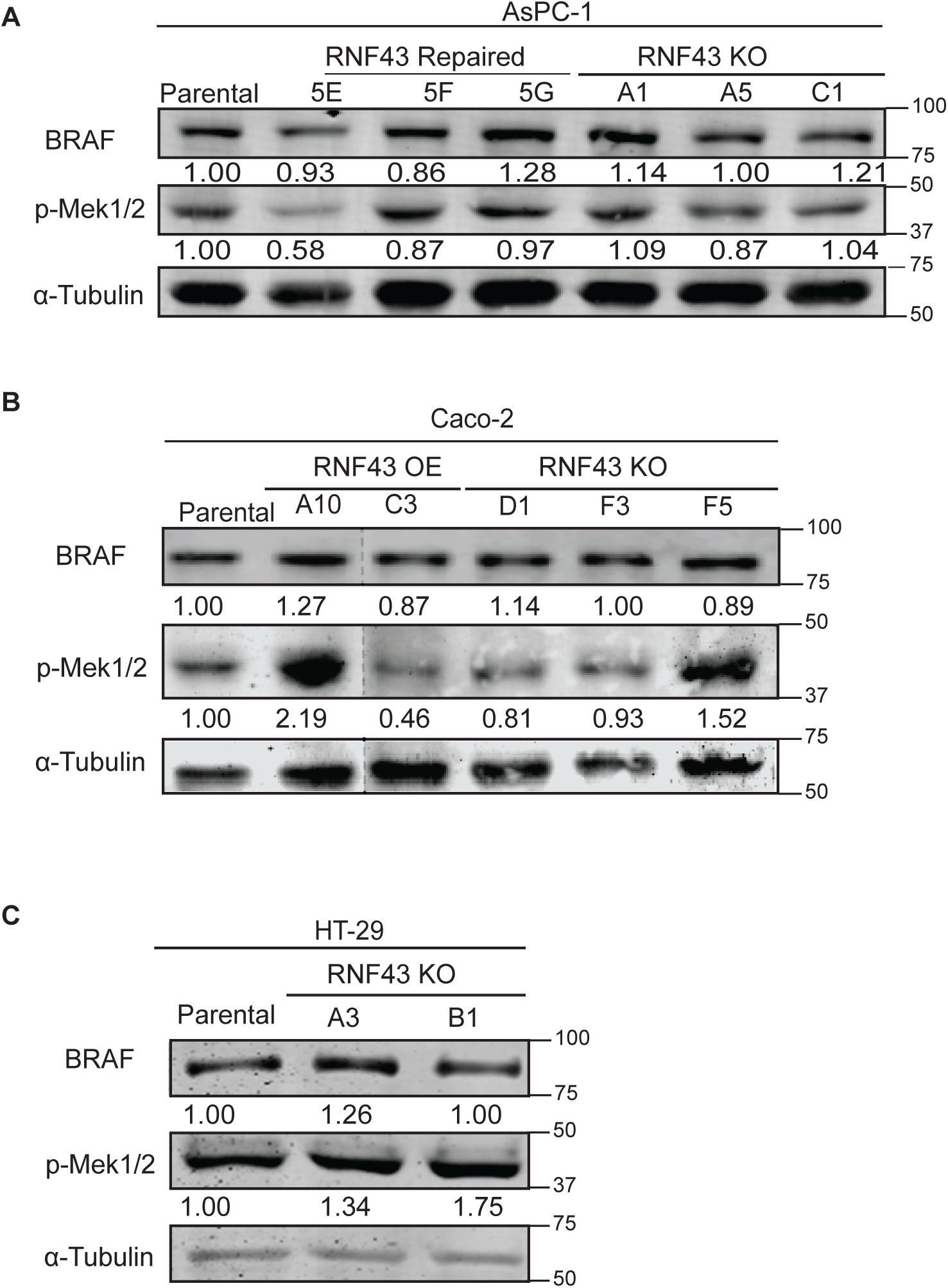
RNF43 status does not correlate with basal BRAF expression or MEK1/2 phosphorylation across pancreatic and colorectal cancer cell lines. (A) Immunoblot analysis of total BRAF and phosphorylated MEK1/2 (p-MEK1/2) in parental AsPC-1 cells, RNF43-repaired clones (5E, 5F, 5G), and RNF43 knockout clones (A1, A5, C1). (B) Immunoblot analysis of total BRAF and p-MEK1/2 in parental Caco-2 cells, RNF43 overexpression (OE) clones (A10, C3), and RNF43 knockout clones (D1, F3, F5). (C) Immunoblot analysis of total BRAF and p-MEK1/2 in parental HT-29 cells and RNF43 knockout clones (A3, B1). α-Tubulin served as a loading control. Numbers below the BRAF and p-MEK1/2 bands indicate relative protein levels normalized to α-tubulin and expressed relative to the parental sample within each panel (set to 1). Data shown are representative of three independent experiments. Where indicated by dashed lines, non-adjacent lanes from the same blot were juxtaposed for presentation.

Under basal conditions, BRAF is primarily localized to intracellular cytosolic compartments, while RNF43 is transferred via the endoplasmic reticulum to a membrane location. As a result, BRAF and RNF43 reside in distinct subcellular compartments, reducing chances for an efficient interaction. However, upon growth factor stimulation, BRAF is translocated to the plasma membrane^[30]^. To test whether this affects RNF43-dependent regulation, cells were serum-starved and subsequently stimulated with HGF and serum for 3 hours. As expected, this treatment increased pMEK1/2 levels and induced a mobility shift of BRAF indicative of multiple phosphorylation events. However, total BRAF levels were essentially unaltered between parental and RNF43-KO HT-29 cells (Supplementary Figure S11B).

Overall, at endogenous expression levels, we find no evidence that RNF43 regulates BRAF, with only minimal effects observed when both proteins are overexpressed. Therefore, we are unable to confirm the findings reported by Hsu et al.^[24]^.

## Discussion

Recently, it was reported that RNF43-mutant colorectal cancers carrying the common BRAF^V600E^ mutation, respond better to a combined BRAF/EGFR inhibitory treatment^[20–22]^. A potential mechanism through which RNF43 loss may have contributed to this improved treatment response, remained however elusive. RNF43, and its homolog ZNRF3, are best known for their role in removing the Wnt-receptor complex from the membrane, thereby keeping Wnt-signaling at minimal essential levels^[1, 3]^. In recent years, both proteins were shown to potentially also regulate other membrane-located proteins as well as some intracellular ones^[3, 5–12, 31]^. Based on these observations, we speculated that RNF43 may also reduce EGFR and/or BRAF protein levels. Theoretically, in RNF43-mutant colorectal cancers, this would lead to increased EGFR and/or BRAF levels, potentially making these cancers more addicted to increased EGFR/BRAF signaling, and indirectly also making them more sensitive to their combined inhibition. Here, we generated several cell line clones in which RNF43 is knocked-out, stably overexpressed, or repaired, to determine if RNF43 can indeed affect EGFR/BRAF levels. We combine these clones with overexpression experiments and R-spondin exposure. Although we occasionally observe a slight clone-to-clone or experimental variation, we did not observe a consistent result indicating that RNF43 can effectively regulate EGFR/BRAF levels, either at endogenous expression or upon overexpression. These results are in contradiction with two recent reports suggesting that this would be the case.

The first report by Yue et al. provides an extensive analysis showing that both RNF43 as well as ZNRF3 can effectively degrade EGFR^[23]^. Similar to our approach, they use various cell line tools in which RNF43 or ZNRF3 expression is knocked-out or overexpressed. In basically all knock-out experiments, just one pool of cells is used in which RNF43/ZNRF3 is “knocked-out” using a lentiviral CRISPR approach. Nevertheless, combined with their co-immunoprecipitation experiments demonstrating a direct interaction between RNF43/ZNRF3 and EGFR via their extracellular domains, and various other experiments, their results appear to firmly demonstrate the claim that RNF43/ZNRF3 regulate membrane and total levels of EGFR. So, why are we unable to confirm these results in our setting?

One potential limitation of the tools generated by Yue et al. is their approach to knockout RNF43 and ZNRF3. Their method relies on the lentiCRISPR-v2 system, in which a lentiviral construct, similar to our pX459 vector, integrates into the genome of target cells and continuously produces both the guide RNA (gRNA) and Cas9, along with the puromycin-resistance protein^[32]^. Due to this sustained expression, the system increases the probability of unintended gene targeting, potentially resulting in off-target effects, some of which could theoretically affect EGFR. A partial solution to this issue is the concurrent generation of a cell pool using a non-targeting lentiCRISPR-v2 vector. However, it is unclear whether this control was consistently applied across all RNF43 knockout pools, especially for the RNF43-KO HT-29 cells that are used for several experiments.

Furthermore, this approach is expected to generate a heterogeneous cell population, where some cells achieve a complete knockout of RNF43/ZNRF3, while others retain either wild-type expression or partial loss of the target gene. Because reliable antibodies for assessing RNF43/ZNRF3 knockout efficiency are unavailable^[33]^, the extent of their expression loss in these ’knockout’ cells remains uncertain. Admittedly, in our own case, direct evaluation of RNF43 protein levels in our knockout clones is also not feasible. However, through our clonal analyses, we can accurately characterize CRISPR-induced DNA modifications, allowing a good evaluation of expected loss of protein expression. Using these clones, we occasionally observed a moderate 1.5-1.75 fold increase in EGFR levels in some of our initial Caco-2 and HT-29 clones.

Importantly, this analysis also showed that increases in EGFR levels were almost exclusively observed in clones that had inadvertently integrated the pX459 Cas9 vector. To assess whether these increases were vector-related rather than RNF43-dependent, we generated multiple Caco-2 and HT-29 clones, varying in pX459 integration status and RNF43 knockout. Surprisingly, EGFR expression did not correlate with RNF43 status, but appeared closely associated with Cas9 integration, suggesting that the modest increases observed in some knockout clones were likely resulting from Cas9 or vector-related effects rather than RNF43 deficiency. Although the underlying mechanism altering EGFR levels remains unclear, and is also beyond the scope of this manuscript, prior studies have reported substantial cellular consequences of Cas9 expression. Sustained Cas9 expression in multiple cell lines was shown to induce DNA damage and activate the p53 pathway, leading to various alterations in transcriptional programs and induction of cellular stress responses^[34–38]^. More concerningly, chromothripsis has been reported as a potential consequence of CRISPR/Cas9-mediated genome editing, underlying the risk of large-scale genomic rearrangements^[39, 40]^. Taken together, these observations indicate that Cas9 itself can have profound effects on cellular phenotypes, including potentially altering EGFR protein levels.

We were also unable to replicate another experiment from Yue et al. demonstrating that overexpressed EGFR can be effectively downregulated by co-expressed RNF43 or ZNRF3 in HEK293T cells. Despite multiple attempts, we fail to observe a consistent decrease in EGFR levels induced by wild-type RNF43 or ZNRF3. In contrast, mature FZD5 levels are downregulated as expected. Notably, some of the authors’ own data hint that this effect may not be fully reproducible, as the input blots for ZNRF3 transfected HEK293T cells in their immunoprecipitation experiments (FiguresLJ4F andLJ5E in their manuscript) also fail to show a clear decrease in EGFR levels.

In theory, R-spondins could influence EGFR levels through two mechanisms. First, by effectively depleting RNF43 and ZNRF3 from the membrane, they would simulate a functional knockout of both proteins, which would be expected to elevate EGFR levels. However, when we treat four different cell lines with Rspo1-conditioned medium or commercial RSPO2, we do not observe the expected rise in EGFR levels. This stands in contrast to the findings of Yue et al., who reported increased EGFR in all cell lines exposed to RSPO2. Second, certain R-spondins have been found to facilitate the formation of a ternary complex with RNF43/ZNRF3 and a membrane-associated protein, resulting in the removal of the latter from the membrane. For instance, BMPR1A is cleared when it simultaneously interacts with RSPO2/3 and RNF43/ZNRF3, while FGFR4 is removed when bound to ZNRF3 and RSPO2^[9–12]^. However, this mechanism does not appear to be active for EGFR, as we do not observe a significant reduction in its levels upon exposure to any RSPO isoform, even in Caco-2 cells overexpressing RNF43. Similarly, EGFR internalization was not affected by any R-spondin variant, in contrast to EGF, which triggers rapid internalization.

Summarizing the EGFR part, we find no evidence that EGFR function is regulated by RNF43/ZNRF3 or R-spondins. This observation is consistent with the findings of Koo et al., who conducted a Mass-Spec analysis to identify membrane proteins that decrease in abundance following RNF43 overexpression^[26]^. EGFR was reported as one of the unaffected proteins. Our results do however strongly contrast those of Yue et al.^[23]^. Given the apparent robustness of several of their results, it is clear that further independent research is needed to establish the definitive relationship between RNF43 and EGFR.

RNF43, ZNRF3, and EGFR all contain N-terminal signal peptides that direct their synthesis to the endoplasmic reticulum, enabling their transport to the plasma membrane. This makes it more plausible for these three proteins to interact and regulate each other. In contrast, BRAF lacks such a signal peptide and is synthesized on free ribosomes, leading to a predominantly cytosolic localization. As a result, at baseline conditions, BRAF and RNF43 reside mostly in distinct subcellular compartments, reducing the likelihood of efficient interaction. This spatial separation makes direct regulation of BRAF protein levels by RNF43 less probable, particularly at endogenous expression levels. Accordingly, we did not observe significant changes in endogenous BRAF or downstream phosphorylated MEK1/2 protein levels in our cell line clones with knock-out, overexpressed or repaired RNF43 expression. BRAF temporarily moves to the membrane when cells are stimulated with growth factors^[30]^, which would allow an interaction with RNF43. We forced this situation by serum-starving HT-29 cells followed by an HGF/FCS chase, but also in this case we did not observe any alteration in BRAF levels.

Our results contrast with a recent study by Hsu et al., in which they indicate that RNF43 effectively targets BRAF for proteasomal degradation through direct interaction and ubiquitination at K499^[24]^. Their analyses primarily depend on transient RNF43 overexpression in mostly pancreatic (cancer) cell lines, leading to a consistent reduction in BRAF protein levels and downstream signaling. However, transient overexpression often results in excessively high expression levels, up to more than a 1000-fold increase compared to endogenous *RNF43* RNA levels in our hands^[25]^. Such high level expression may lead to more frequent intracellular encounters of RNF43 and BRAF, thereby possibly promoting BRAF degradation, while this would infrequently occur at endogenous levels.

Further support for their findings is provided by two CRISPR/Cas9 *RNF43* gene-edited pancreatic cell lines. While their methods section states that single-cell derived clones were generated, no details are provided about the specific DNA alterations induced that may lead to RNF43 protein loss. A single immunoblot is presented, showing reduced RNF43 levels rather than a complete loss of expression, which correlates with increased BRAF levels, supporting their findings. However, researchers in this field are well aware that no RNF43 antibody is currently available that reliably detects endogenous RNF43 expression^[33]^. As a result, it remains uncertain whether the observed bands genuinely represent RNF43 or if they correspond to another protein of similar size detected nonspecifically.

Taken together, the reliance on RNF43 overexpression and the uncertainty surrounding their RNF43 knock-out cell line models, raise doubts about whether RNF43-mediated BRAF downregulation is relevant under endogenous expression conditions. This skepticism is reinforced by our own analysis, in which we fail to identify any correlation between endogenous RNF43 function and BRAF levels or downstream signaling.

Over the past decade, several potential substrates of RNF43 and ZNRF3 have been proposed^[3]^. Our findings, however, suggest that neither EGFR nor BRAF are genuine targets of RNF43LJmediated degradation. Because this differs from two recent reports, further independent validation will be important to clarify how these proteins relate to RNF43. Together with the now-retracted report that had implicated E-cadherin as an RNF43 substrate^[41, 42]^, our observations indicate that the spectrum of proteins regulated by RNF43/ZNRF3 may be narrower than previously assumed and more confined to components of the Wnt signaling pathway. Furthermore, we initiated our research by speculating that RNF43/BRAF^V600E^ mutant colorectal cancers may respond better to a combined BRAF/EGFR inhibition, because of increased EGFR and BRAF levels following RNF43 loss. As our findings did not reveal a consistent connection between RNF43 function and the regulation of EGFR or BRAF levels, other explanations will need to be explored. Finally, we observed that stable integration of the pX459 gene-editing vector can aberrantly elevate EGFR expression, underscoring the need for caution when using cell line models with persistent Cas9 expression in research and clinical settings.

## Methods

### Cell lines and culture

HEK293T (RRID: CVCL_0454), Caco-2 (RRID: CVCL_0025), HT-29 (RRID: CVCL_A8EZ), and HeLa (RRID: CVCL_0030) cells were cultured in Dulbecco’s Modified Eagle medium (DMEM; Lonza, Breda, the Netherlands) supplemented with 10% fetal bovine serum (FBS; Gibco, Bleiswijk, the Netherlands). AsPC-1 (RRID:CVCL_0152) cells were maintained in Roswell Park Memorial Institute (RPMI)-1640 culture medium (Lonza), also supplemented with 10% FBS. Media were refreshed every 2–3 days. All cell lines were cultured in a humidified incubator maintained at 37 °C with 5% CO_2_. The identity of all cell lines and clones was confirmed by the Erasmus Molecular Diagnostics Department, using Powerplex-16 STR genotyping (Promega, Leiden, the Netherlands). Mycoplasma contamination was ruled out in all lines through a real-time PCR method by Eurofins GATC-Biotech (Eurofins GATC Biotech, Konstanz, Germany).

### Plasmids used in this study

A C-terminal 3×FLAG-tagged RNF43 expression construct designed for stable integration was generated by first replacing the original CMV promoter with the EF-1α promoter using Gibson assembly (New England Biolabs, Ipswich, MA, USA) ^[43]^. Next, Gibson assembly was used to replace the original Puromycin selection marker with Hygromycin. This backbone was used to generate C-terminal HiBiT-tagged R-spondin vectors, in which the 3xFLAG tag was replaced with the HiBiT tag preceded by a SGGGSGGGSG linker sequence using Q5 mutagenesis (New England Biolabs). Vectors containing the human *RSPO1*, *RSPO2* and *RSPO3* open reading frames were kindly provided by Dr. Niehrs from the DKFZ, Heidelberg, Germany. The RSPO1, RSPO2 and RSPO3 fragments were separately assembled into the C-terminal HiBiT-tagged vector using Gibson Assembly. Full-length cDNA of human *RSPO4* was acquired from the human esophageal cancer TE8 cell line by RNA purification using the NucleoSpin RNA kit (Macherey-Nagel, Düren, Germany), followed by reverse transcription PCR with specific primers and the high-capacity cDNA reverse transcription kit (Applied Biosystems, Foster City, CA, USA). The ORFs of both long and short isoform of RSPO4 were assembled into the C-terminal HiBiT-tagged RNF43 vector with EF-1α promoter using Gibson Assembly.

The C-terminal FLAG-tagged RNF43 wild-type and RING domain variant (W302R) and the C-terminal 3xFLAG-tagged ZNRF3 wild-type and RING domain variant (H292Y) were generated previously^[44]^. C-terminal HA-tagged RNF43 and ZNRF3 expression vectors were also available^[44]^.

The EGFR-GFP construct was kindly provided by Alexander Sorkin (Addgene plasmid # 32751). We also generated an EGFR expression plasmid in which the GFP tag was removed using Q5 Site-Directed Mutagenesis. The p3xFLAG CMV14/BRAF-WT and BRAF-V600E plasmids were a gift from Jacques De Grève (Addgene plasmid # 131710 and 131723). The V5-tagged Frizzled5 (FZD5) construct was generously provided by Dr. Maurice (UMC Utrecht, The Netherlands).

Cloned sequences of all plasmids were verified for full-length sequence integrity. The sequences of all primers used in this study are provided in Supplementary Table S1.

### Generation of conditioned medium

To generate L-RSPO conditioned media (RSPOs-CM), mouse fibroblast L-cells were transfected with RSPO-HiBiT plasmids driven by the EF-1α promoter. Clones in which the constructs were stably integrated were selected by Hygromycin (600 μg/mL) selection for 1 week, followed by picking of single clones. Clones expressing adequate amounts of each R-spondin were selected by a HiBiT Assay with the Nano-Glo HiBiT Extracellular Detection System (Promega) (Supplementary Figure S9A). Correct expression of each R-spondin variant was confirmed with immunoblotting using a anti-HiBiT antibody according to standard procedures (Supplementary Figure S9B). RSPO1-conditioned medium was collected from HEK293T cells stably expressing mouse RSPO1. Human recombinant R-spondin 1 (PeproTech, #120-38), R-spondin 2 (PeproTech, #120-43), and R-spondin 3 (PeproTech, #120-44) were used as controls (5 μg/mL). Functionality of each R-spondin variant was tested in a β-catenin reporter assay (Supplementary Figure S9C).

Next, all L-RSPOs variant clones were seeded at ∼10% density in 100-mm dishes with DMEM and 10% FBS, cultured for 4 days, and the conditioned media were collected. Fresh medium was added, and cells were incubated for another 3 days. The media were then filtered and stored at -20°C, and were used at a 1:10 dilution.

For Wnt3a treatment, cells were cultured in 10% L-Wnt3A-conditioned medium (Wnt3A-CM) or L-control-conditioned medium (Control-CM) according to standard procedures^[44]^.

### Quantitative PCR

Total RNA was extracted with the NucleoSpin RNA kit (Macherey-Nagel) and reverse-transcribed using the PrimeScript RT Reagent Kit (TaKaRa) following the manufacturer’s guidelines. Quantitative PCR was conducted on the StepOne Real-Time PCR System (Applied Biosystems) in triplicate. Gene expression levels were analyzed using the ΔΔCT method, normalizing to *GAPDH*. Primer sequences are provided in Supplementary Table S1.

### Luciferase reporter assays

Wnt-Responsive Element (WRE) β-catenin luciferase assays were carried out as previously reported^[25, 45]^. Briefly, HEK293T or AsPC-1 cells were transiently transfected using Lipofectamine 2000 (Life Technologies). After 6 hours, cells were stimulated with RSPO-CM, Wnt3A-CM or L-Control conditioned medium, and cultured for an additional two days. Firefly and Renilla luciferase activity was measured with the Dual-Luciferase Reporter Assay system (Promega) according to the manufacturer’s instructions in a LumiStar Optima luminescence counter (BMG LabTech, Offenburg, Germany). Firefly luminescence was normalized to Renilla. Experiments were performed in triplicate.

### Western blots

For western blot experiments, cells were washed twice with PBS and lysed in 2x Laemmli buffer supplemented with 0.1 M DTT. Subsequently, lysate<u>s</u> were boiled for 10 minutes at 95°C. For immunoblotting, membranes were blocked with 5% nonfat Dry Milk (Cell Signaling Technology, Danvers, Massachusetts, USA) or Odyssey blocking buffer (LI-COR Biosciences, Lincoln, NE, USA). For fluorescent immunoblotting, proteins were detected on the Odyssey Infrared Imaging system (LI-COR Bioscience). For enhanced chemiluminescence (ECL)-based detection, Immobilon Block-CH (Chemiluminescent Blocker) blocking buffer was used (cat. #WBAVDCH01, Millipore). The secondary antibody used was Goat anti-rabbit/HRP (dilution 1:3000, catalog #7074, Cell Signaling Technology). Membranes were then incubated with Immobilon ECL Ultra Western HRP Substrate (Millipore) for 3-5 minutes and visualized using the Amersham Imager 600 (GE Healthcare). Details of the antibodies used in this study can be found in Supplementary Table 2. All full western blot images can be found in Supplementary Figure S12.

### CRISPR/Cas9 genome editing and clone screening

RNF43 knockout in AsPC-1, Caco-2 and HT-29 cells was generated via CRISPR/Cas9 genome editing. Guide RNAs (gRNAs) (Supplementary Figure S2), which were designed with the help of the CRISPOR (crispor.tefor.net), were cloned into the pSpCas9(BB)-2A-GFP (pX458, Addgene #48138) or pSpCas9(BB)-2A-Puro (pX459, Addgene #48139) vector, gifts from Feng Zhang, using standard procedures^[46]^. Cells were seeded into a 6-well plate and transfected with 2.5 μg of pX458 or pX459 using Lipofectamine 3000, following the manufacturer’s instructions. For AsPC-1, GFP-positive cells were sorted out 48 hours following transfection, and plated into 96-well plates by a FACSAria II cell sorter (BD Biosciences). Caco-2 and HT-29 cells were trypsinized to 10 cm petri dishes. Caco-2 cells were treated with 3 μg/ml puromycin for 2 days, while HT-29 cells were treated with 2 μg/ml puromycin for 2 days. Clones that appeared were hand-picked and transferred to 96-well plates.

The RNF43 S720* mutation present in Aspc-1 cells was repaired using CRISPR/Cas9 technology. To achieve this, a 1579 bp genomic PCR fragment from WT DNA was cloned into the pGEM-T Easy Vector (cat. #A1360, Promega), serving as a homology-directed repair (HDR) template. Primers can be found in Supplementary Table S1. Silent mutations were introduced using Q5 mutagenesis to prevent Cas9 re-cutting after successful repair (see Supplementary Figure S4). In addition, a sgRNA was cloned into pSpCas9(BB)-2A-GFP (pX458). Transfection and sorting were performed as described above.

To generate HT-29 panels differing in Cas9 status and RNF43 genotype, we transfected cells with empty pX459 or carrying an RNF43-targeting gRNA, as described above. RNF43 WT clones were obtained by transfection with empty pX459 followed by a transient 2-day puromycin selection (2 μg/mL) and single-colony isolation. RNF43 knockout clones were generated by transfection with pX459-RNF43 gRNA followed by identical selection and clonal isolation. After expansion, RNF43 editing was confirmed by Sanger sequencing (Supplementary Figure S12), and clones were stratified by Cas9 status based on Cas9 detectability by immunoblotting, yielding Cas9-positive and Cas9-negative clones within both RNF43 WT and RNF43 KO backgrounds. To obtain Caco-2 RNF43 WT clones with different Cas9 status, Caco-2 cells were transfected with empty pX459, puromycin selected (3 μg/ml puromycin for 2 days), and single colonies were isolated and subsequently classified as Cas9-positive or Cas9-negative using the same immunoblot-based criterion.

Two weeks later, both genomic DNA and cDNAs were generated from the expanded single-cell clones. Genomic DNA was extracted using QuickExtract DNA Extraction Solution (Epicenter, Madison, WI, USA), while cDNA was obtained as previously described. To verify that the correct RNF43 alterations were obtained, PCR fragments encompassing the sgRNA site were generated, followed by Sanger sequencing (Macrogen). The sequence alterations and the primers used for RNF43 verification are shown in Supplementary Figure S2.

### Generation of Caco-2 RNF43 overexpression clones

Caco-2 cells were seeded into a 6-well plate and transfected with 500 ng of EF1α-RNF43-3xFLAG plasmid using Lipofectamine 2000, following the manufacturer’s instructions. 48 hours post-transfection, the cells were subjected to selection with 600 µg/mL hygromycin for 10 days. Emerging clones were manually picked and transferred to 96-well plates for further expansion.

Two weeks later, both protein and cDNA were collected from the expanded single-cell clones. cDNA was obtained as previously described. To confirm the successful introduction of the exogenous RNF43, RT-qPCR and Western blot analyses were performed.

### EGFR Immunofluorescence analysis

AsPC-1 and Caco-2 cells were grown on coverslips in 12-well plates in culture medium without phenol red. After treating with EGF (GenScript, Cat.#Z00333) or RSPOs for 30 minutes or 2 days, cells were washed with PBS three times. Then, cells were fixed with 4% PBS-buffered formaldehyde solution at room temperature for 10LJminutes. After PBS washing three times, cells were permeabilized with 0.2% Triton X-100 in PBS for 10 min, washed with PBS, and incubated with blocking solution (BSA 3% in PBS) for 30LJminutes at room temperature. Cells were then stained with anti-EGFR antibody (1:500, Cell Signaling Technology, cat.# 54359) overnight at 4 °C. After three times washing with PBS, cells were incubated for 1 h at room temperature with donkey anti-Rabbit-Alexa 488 (Thermo Fisher Scientific, cat.# A-21206) protected from light. Next, cells were mounted in VECTASHIELD antifade mounting medium with DAPI (cat. # H-1200, Vector Laboratories, Burlingame, CA, USA). Images were captured by a Zeiss LSM700 confocal laser scanning microscope using ZEN 2009 software with constant parameter setting.

### FACS analysis

Cells were trypsinized and resuspended in antibody dilution buffer (0.5% BSA in PBS). A total of 5 × 10^5^ cells were incubated with the EGFR (D1D4J) XP Rabbit antibody (PE Conjugate) (Cell Signaling Technology, cat.# 48685) for 1 hour at 4 °C. Following incubation, the cells were washed three times with the antibody dilution buffer and then resuspended in the same buffer. Flow cytometry was conducted using a BD FACS Canto II (BD Biosciences, CA, USA), and the resulting data were analyzed with FlowJo software.

### Statistical analysis

Data are presented as mean ± SD. Statistical analyses were performed using GraphPad Prism v8.0.2 (GraphPad Software, San Diego, CA, USA; RRID: SCR_002798). A two-sided P value < 0.05 was considered statistically significant. Significance is indicated as n.s. (not significant), *P < 0.05, **P < 0.01, ***P < 0.001, and ****P < 0.0001.

### Author contributions

JN: Generation and interpretation of data, writing the manuscript, SL, RZ, JM: Generation and interpretation of data, MPP: Supervision of the project, RS: Conceptualization and hypothesis, data generation and interpretation, writing the manuscript. All authors reviewed the results and approved the final version of the manuscript.

### Data availability statement

All data supporting the findings of this study are available within the paper and its Supplementary Information.

## Supporting information

Suppmenentary Figures

