## Supplementary material for "Reassessment of RNF43 Function Reveals No Impact on Endogenous EGFR or BRAF Protein Stability": Suppmenentary Figures

**Supplementary Fig S1.** A newly discovered heterozygous E62* mutation in *ZNRF3* in the AsPC-1 cell line.

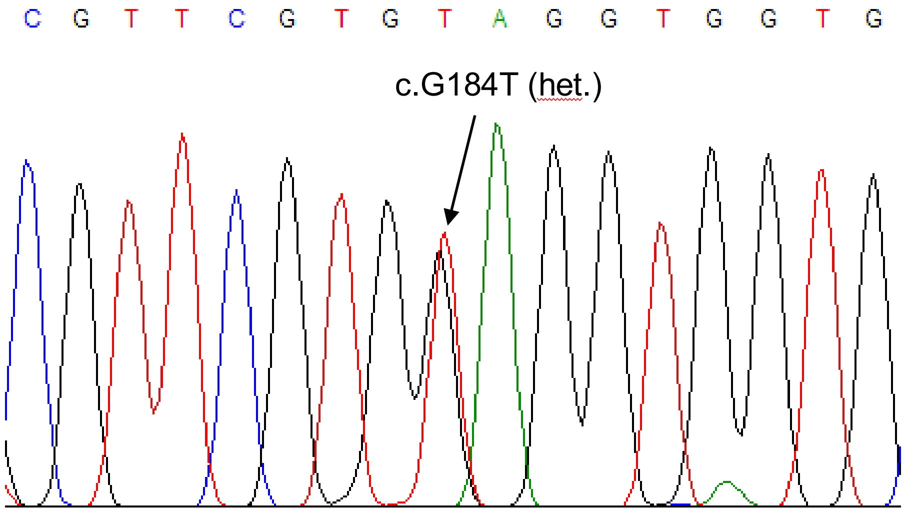

**Supplemental Figure S2.** Depiction of first batch of CRISPR/Cas9 induced RNF43 knock-out mutations.

| Exon 2 as the on-target locus. | | | |
|---|---|---|---|
| Target cell Lines | Oligo name | Oligo sequence | Guide sequence with PAM |
| AsPc-1 | hRNF43-crF2 | CACCgAGGCTTTGGACGCACAGGAC | AGGCTTTGGACGCACAGGAC TGG |
| | hRNF43-crR2 | AAACGTCCTGTGCGTCCAAAGCCTc | |
| Caco-2<br>HT-29 | RNF43-grna4-exon2-Fw | CACCgCAGGCAGGCTTTGGACGCAC | CAGGCAGGCTTTGGACGCAC AGG |
| | RNF43-grna4-exon2-Rv | AAACGTGCGTCCAAAGCCTGCCTGc | |

**AsPC-1**

Clone A1:

CTGCAGGCAGGCTTTGGACGCACAGGACTGGTACTGGCAGCAGCGGTGG (Parental)
CTGCAGGCAGGCTTTGGACGCACAG-ACTGGTACTGGCAGCAGCGGTGG (-1bp) p.G29Dfs*21

Clone A5:

GGACGCACAGGACTGGTACTGGCAGCAGCGGTGG-3' (Parental)
GACGCACAGCCTGGCCCAGTGAGCAAGGGCGAGGAGCTGTTCACCGGGGTGGTGCCCATCCTGGTCGAGC
TGGACGGCGACGTAAACGGGACTGGTACTGGCAGCAGCGGTGG (+80bp) p.G29Afs*48

Clone C1:

GGACGCACAGGACTGGTACTGGCAGCAGCGGTGGAGTCTGAAAGAT (Parental)
GGACGCACA-GACTGGTACTGGCAGCAGCGGTGGAGTCTGAAAGAT (-1bp) p.G29Dfs*21
GGACGCACA-------------------------------AAAGAT (-31bp) p.G29Kfs*11

**Caco-2**

Clone D1:

ACCCTGCAGGCAGGCTTTGGACGCACAGGACTGGTACTGGCA (Parental)
ACCCTGCAGGCAGGCTTTGGAC----------GGTACTGGCA (-10bp) p.T28Yfs*19

Clone F3:

GGCCCTGGCTGCTGATGGCTACCCTGCAGGCAGGCTTTGGAC (Parental)
TACCCTGCAGGCAGGCTTTGGACG----GGACTGGTACTGGCAGCAGC (-4bp) p.T28Dfs*21
GGCCCTGGC----------------------------------------------------
AGCAGCGGTGGAGTCTGAAAGATCAGCAGA (-52bp) p.L16Qfs*17

Clone F5:

CTTTGGACGCACAGGACTGGTACTGGCAGCAGCGGTG (Parental)
CTTTGGAC-CACAGGACTGGTACTGGCAGCAGCGGTG (-1bp) p.T28Qfs*22

**Supplemental Figure S2**. Continued.

**HT-29**

Clone A3:

CCCTGGCTGCTGATGGCTACCCTGCAGGCAGGCTTTGGACGCACAGGACTGGT(Parental)
CCCTGG-----------------------------------------ACTGGT(-41bp)p.L16Tfs*9
CCCTGGCTGCTGATGGCTACCCTGCAGGCAGGCTTTGGAC-----GGACTGGT(-5bp) p.L30Tfs*7

Clone B1:

TGGTGGCCACCAGCTGCAGCTGGCTGCCCTCTGGCCCTGGCTGCTGATGGCTACCCTGCAGGCAGGCTTT
GGACGCACAGGACTGGTACTGG (Parental)
TGGTGGCC----TGCCTGCAGGGTAGCCATCAGCAGCCAGGGCCAGAGGGCAGCCAGCTGCAGC------
-----------------------------GGTGGAGTCTGAAAGA
(52bp inversion and 39bp deletion)p.H5Lfs*4

Fig S3.

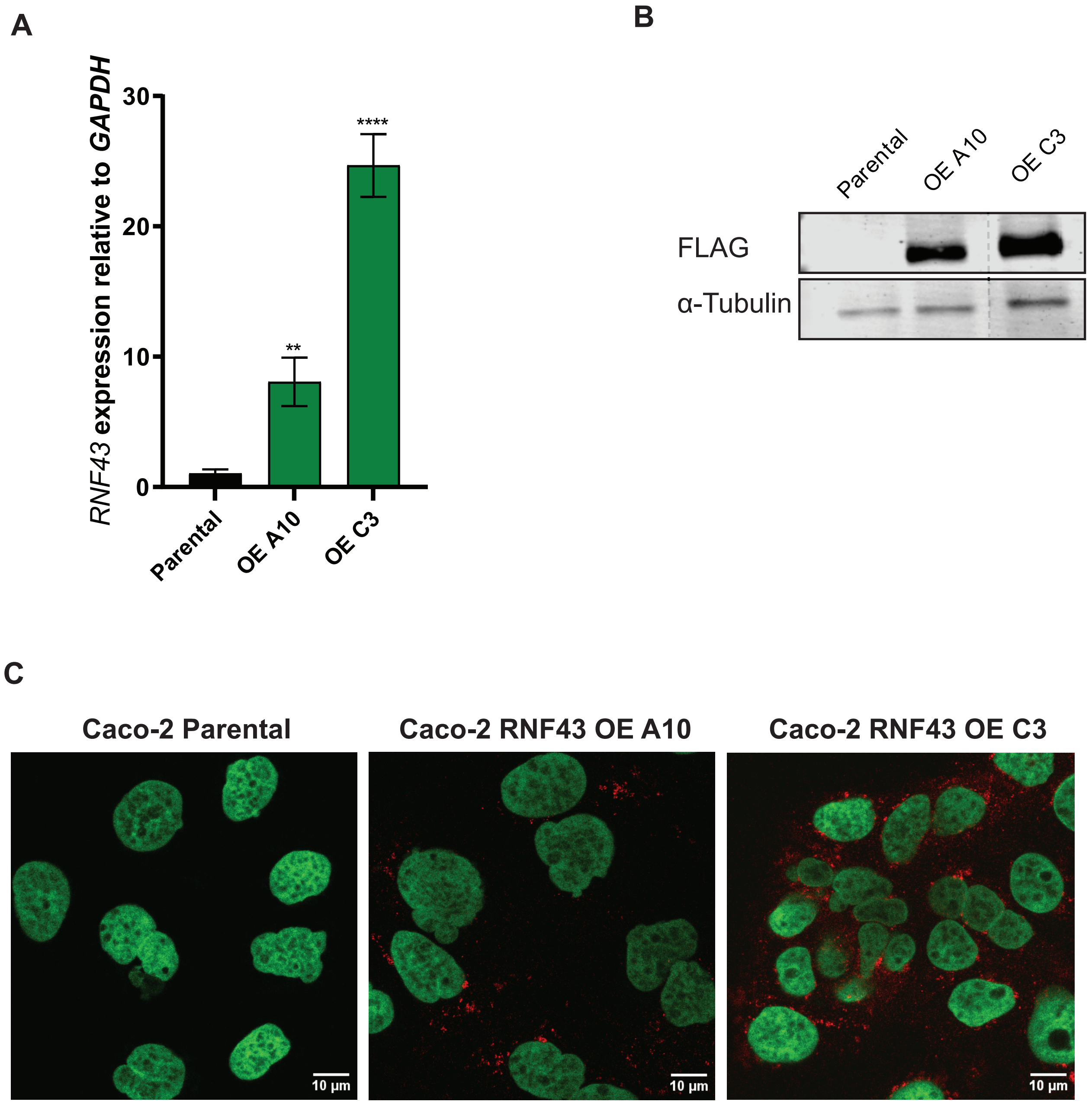


**Supplementary Fig S3.** Generation of Caco-2 clones stably overexpressing FLAG-tagged RNF43.

(A) qRT-PCR analysis showing that clones A10 and C3 express RNF43 at approximately 8-fold and 25-fold higher levels, respectively, compared to endogenous RNF43.

(B) Immunoblot demonstrating correct expression of FLAG-tagged RNF43.

(C) Immunofluorescence using anti-FLAG antibody (red) to visualize the subcellular localization of overexpressed RNF43.

OE, overexpression.

**Supplemental Figure S4.** Sequence of gRNA and repair plasmid to restore original open-reading frame of RNF43 in AsPC-1 cells.

Wild-type RNF43 sequence around codon 720 (underlined):

5'-CCAGAAACCCCAGGCCCCTGTTACTCAAATTCACAGCCAGTGTGGTTGTGC-3'
3'-GGTCTTTGGGGTCCGGGGACAATGACTTTAAGTGTCGGTCACACCAACACG-5'
P E T P G P C Y S N S Q P V W L C

S720* RNF43 mutation present in AsPC-1:

Stop codon in bold letter type; gRNA marked light-blue; PAM-sequence underlined.

5'-CCAGAAACCCCAGGCCCCTGTTAC**TGA**AATTCACAGCCAGTGTGGTTGTGC-3'
3'-GGTCTTTGGGGTCCGGGGACAATGACTTTAAGTGTCGGTCACACCAACACG-5'
P E T P G P C Y *

Sequence as present in repair plasmid:

Silent mutations (red letters) introduced in repair plasmid to prevent recutting by Cas9 following successful repair.

5'-CCAGAAACCCCAGG**A**CCCTG**C**TA**T**TCAAATTCACAGCCAGTGTGGTTGTGC-3'
3'-GGTCTTTGGGGTCCTGGGACGATAACTTTAAGTGTCGGTCACACCAACACG-5'
P E T P G P C Y S N S Q P V W L C

Fig S5.

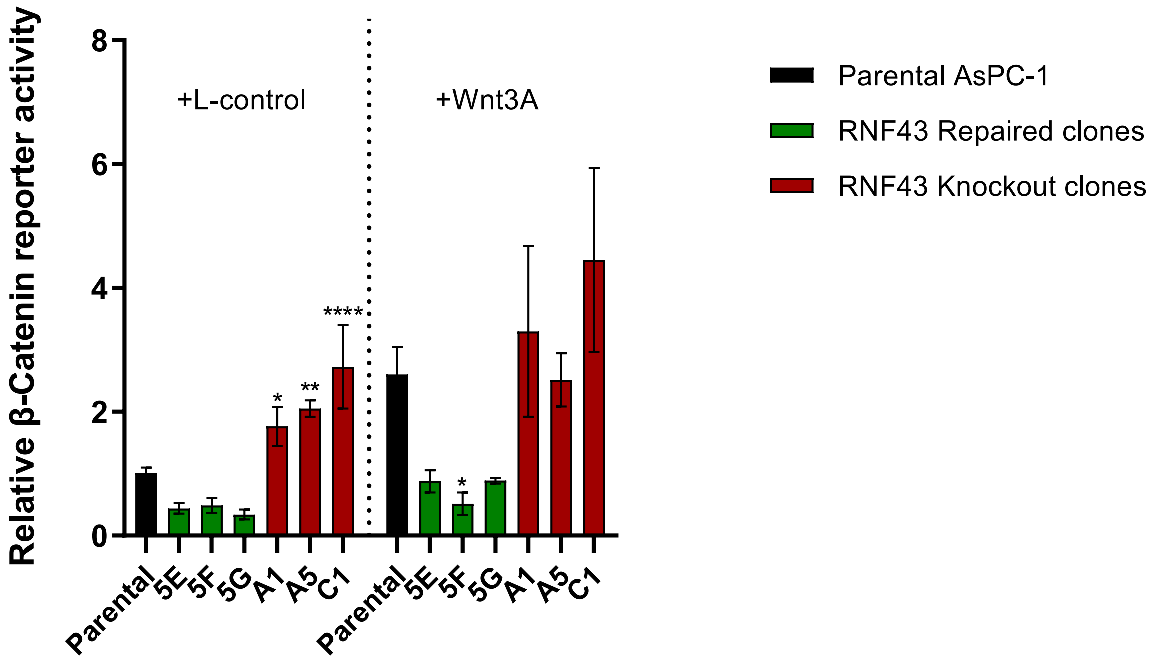


**Supplementary Fig S5.** A β-catenin reporter assay performed on parental, RNF43-repaired, and RNF43 knockout AsPC-1 clones. Parental AsPC-1 cells, RNF43-repaired clones, and RNF43 knockout clones were treated with L-control or L-Wnt3A conditioned medium. Reporter activity is shown as the ratio of WRE-driven Firefly luciferase to CMV-driven Renilla luciferase and is normalized to parental cells under L-control (set to 1). Data are presented as mean ± SD. Statistical significance was assessed by one-way ANOVA with 3 multiple-comparisons test; *P < 0.05, **P < 0.01, ***P < 0.001, ****P < 0.0001.

Fig S6.

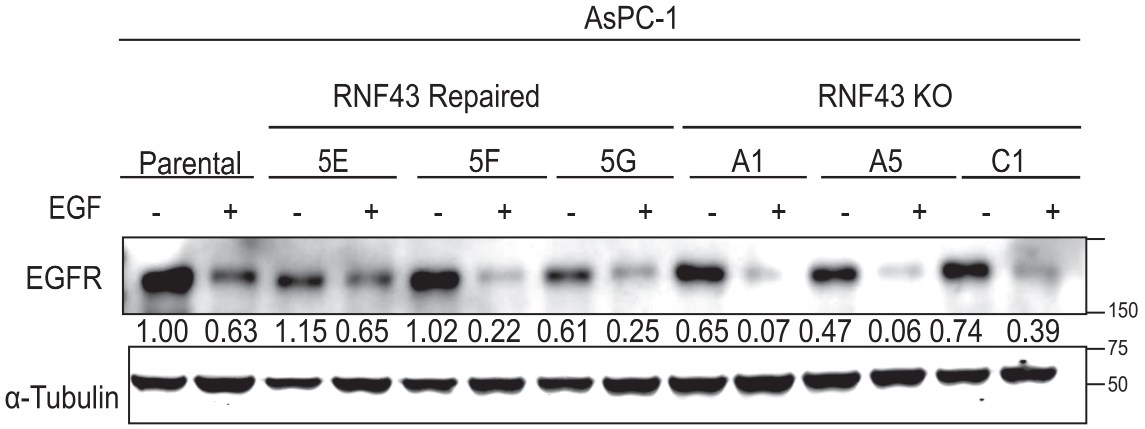


**Supplementary Fig S6.** Parental AsPC-1 cells, RNF43-repaired clones (5E, 5F, 5G), and RNF43 knockout clones (A1, A5, C1) were treated with EGF (5 ng/mL; +) or medium alone (−) for 48 h (culture conditions as described in Methods). Whole-cell lysates were immunoblotted for EGFR; α-tubulin served as a loading control. Numbers indicate EGFR band intensities normalized to α-tubulin and expressed relative to the parental, −EGF condition (set to 1).

Fig S7.

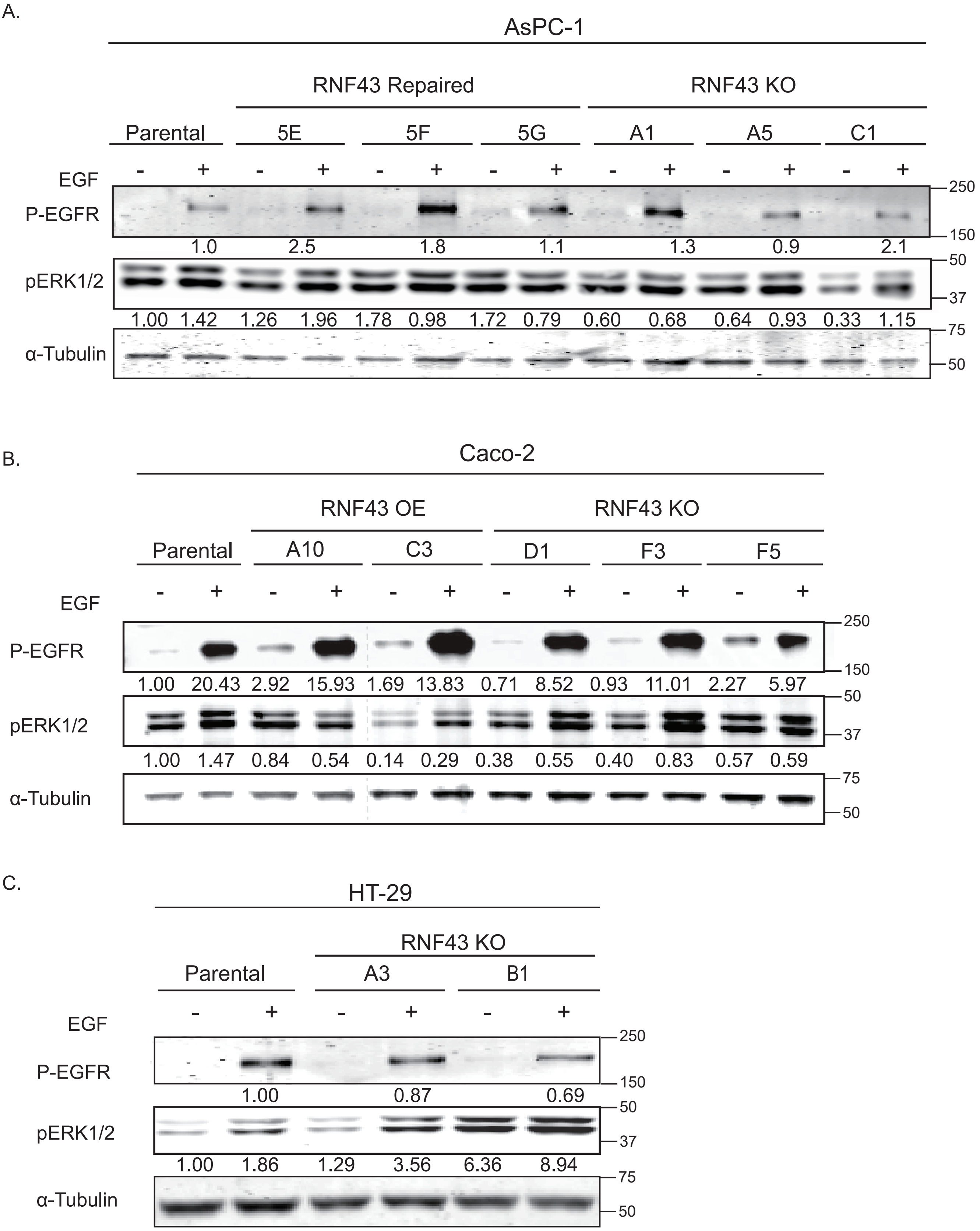


**Supplementary Fig S7.** Analysis of phosphorylated EGFR and ERK1/2, following EGF addition to clones of (A) AsPC-1, (B) Caco-2, and (C) HT-29. Cells were treated with EGF (+) or left untreated (−) for 30 min (EGF, 5 ng/mL; culture conditions as described in Methods). Whole-cell lysates were immunoblotted for phospho-EGFR (p-EGFR) and phospho-ERK1/2 (p-ERK1/2). α-Tubulin served as a loading control. Numbers indicate band intensities normalized to α-tubulin and expressed relative to the parental, −EGF condition (set to 1). Despite clonal variability, p-EGFR and p-ERK1/2 levels do not show a consistent trend across RNF43 genotypes. Where indicated by dashed lines, lanes were juxtaposed from non-adjacent positions on the same blot for presentation.

Fig S8.

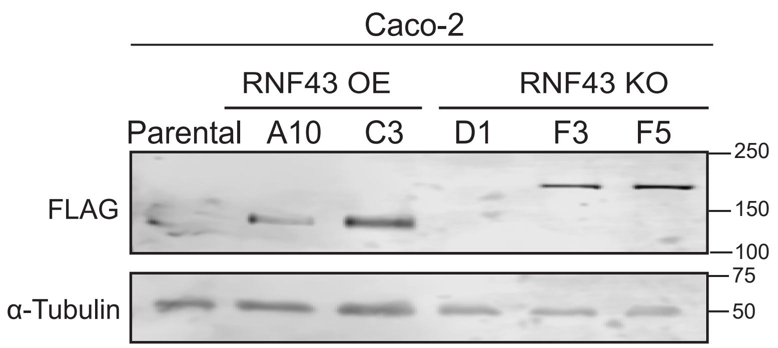


**Supplementary Fig S8.** Immunoblot confirming expression of FLAG-tagged RNF43 (~86 kDa) in Caco-2 RNF43 OE clones, and unintended expression of FLAG-tagged Cas9 in 2 out 3 Caco-2 RNF43 knockout clones. α-Tubulin served as a loading control.

**Fig S9.** Generation of L-cell clones stably expressing C-terminal HiBiT-tagged R-spondin variants.

(A) Secreted R-spondin levels were quantified using a Nano-Glo HiBiT extracellular assay. The left panel shows results from transiently transfected L-cells, and the right panel shows results from stable clones expressing the same variants.

(B) Immunoblot showing the expression of R-spondin variants using an anti-HiBiT tag antibody.

(C) β-catenin reporter assay in HEK293T cells treated with L-Wnt3A conditioned medium together with the indicated R-spondin variants. Conditioned media were obtained from L-cell clones stably expressing the indicated R-spondins and were used at a 1:10 dilution. Recombinant human RSPO1, RSPO2, and RSPO3 (PeproTech) were used at 5 µg/mL (final concentration). Reporter activity is shown as the ratio of WRE-driven Firefly luciferase to CMV-driven Renilla luciferase and is normalized to HEK293T cells treated with Wnt3A alone (set to 1). Data are presented as mean ± SD. ****P < 0.0001; n.s., not significant.

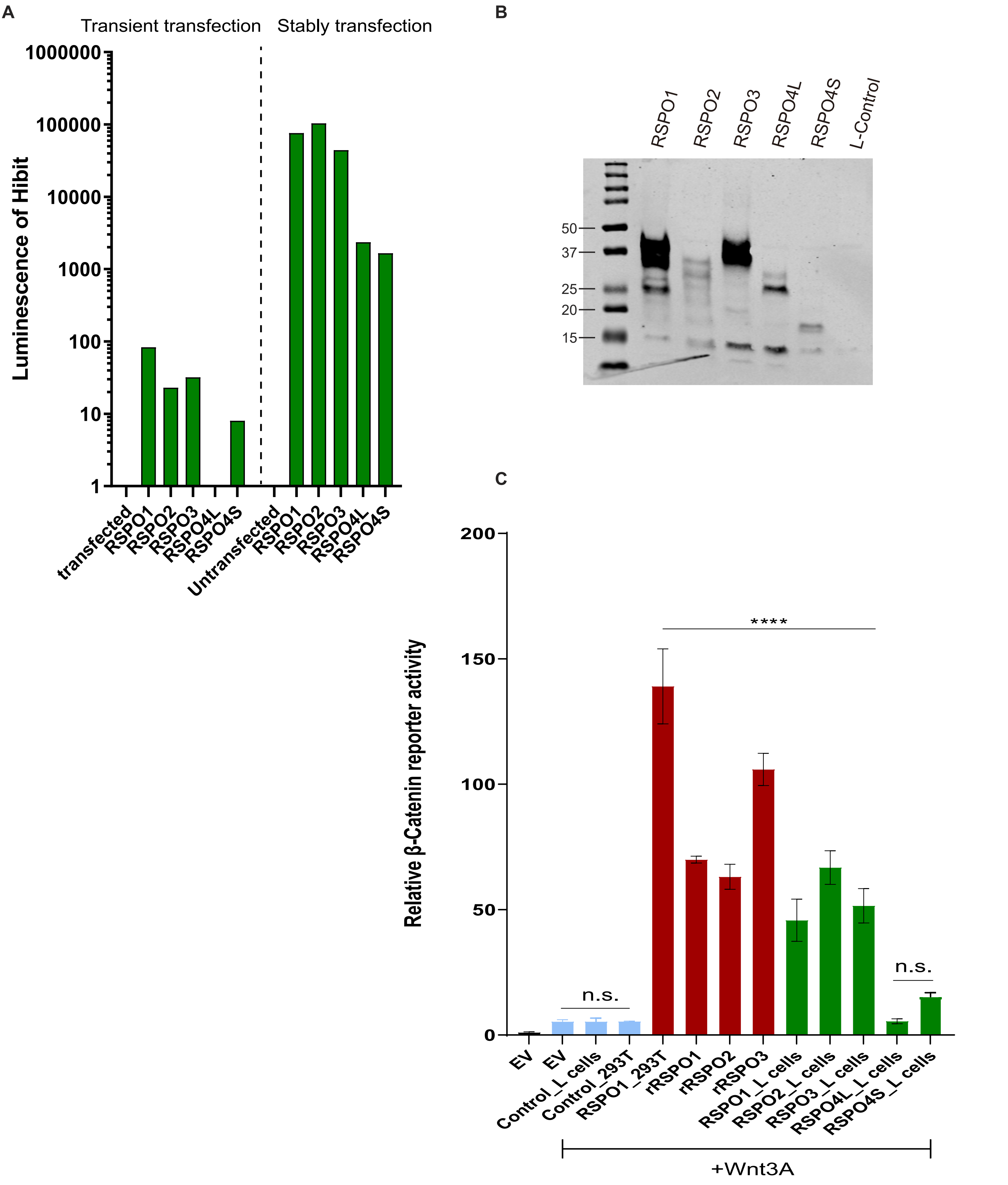

Fig S10.

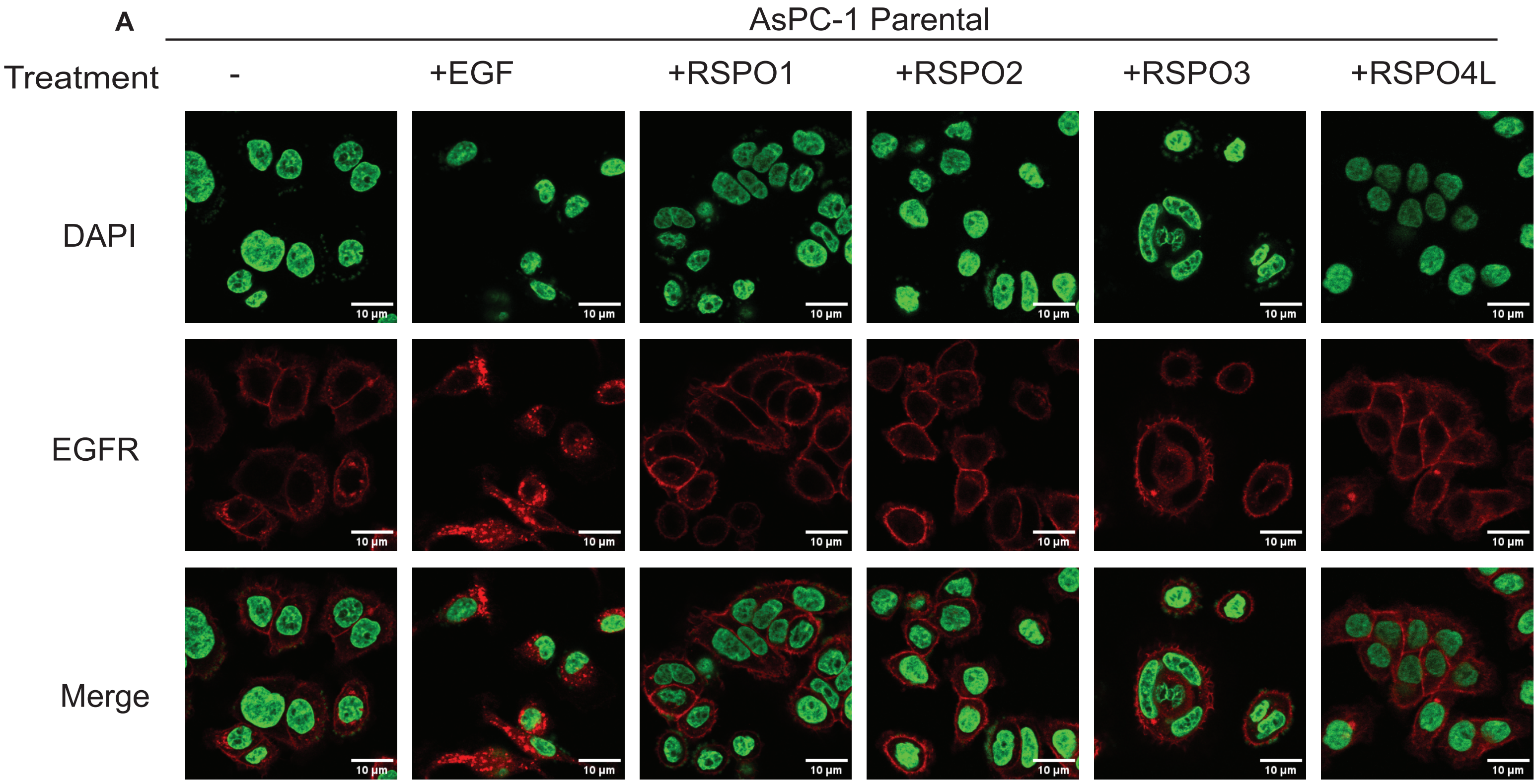
A
AsPC-1 Parental
Treatment
-
+EGF
+RSPO1
+RSPO2
+RSPO3
+RSPO4L
DAPI
EGFR
Merge
10 µm

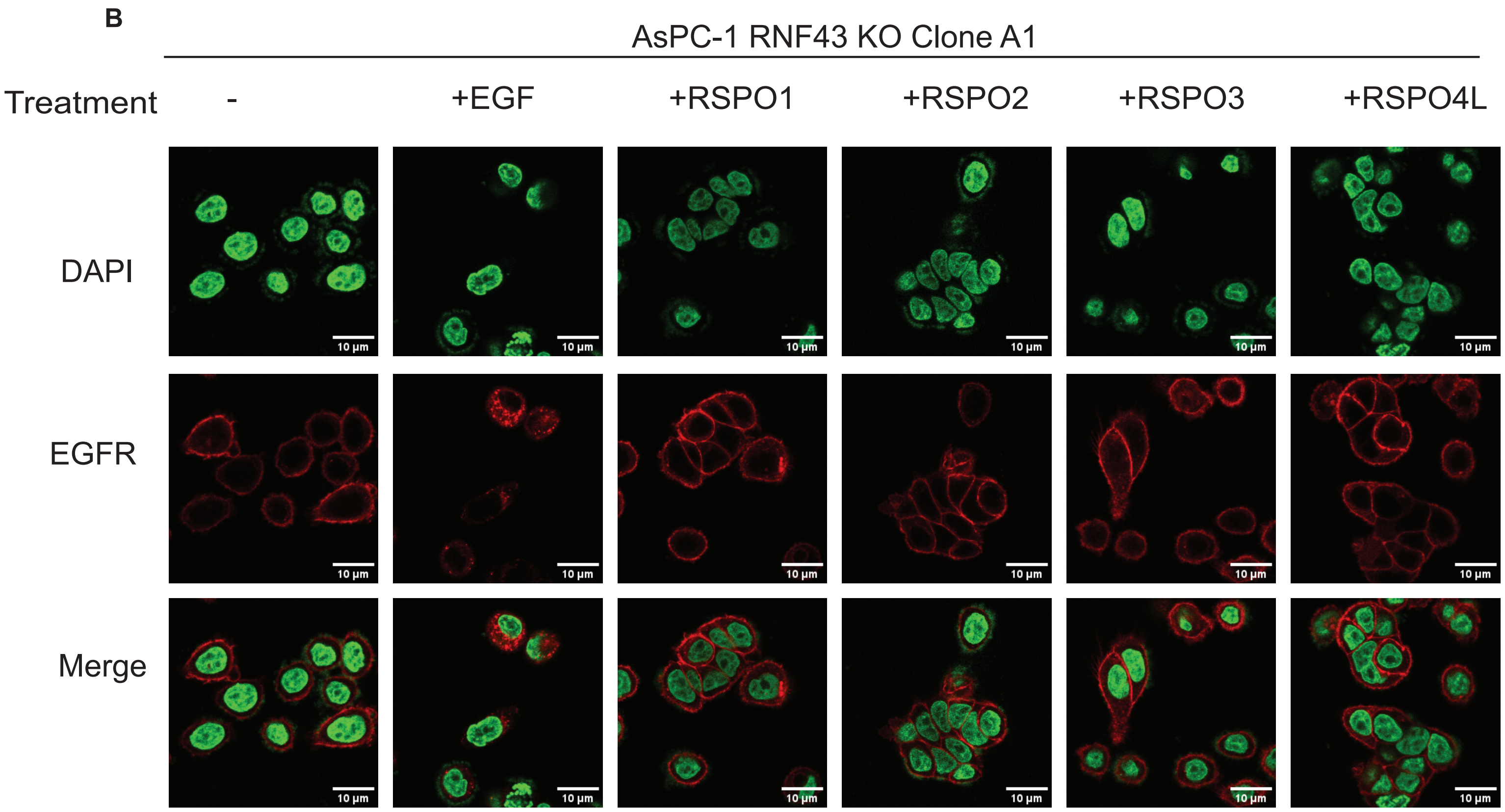
B
AsPC-1 RNF43 KO Clone A1
Treatment
-
+EGF
+RSPO1
+RSPO2
+RSPO3
+RSPO4L
DAPI
EGFR
Merge
10 µm

Fig S10. Continued

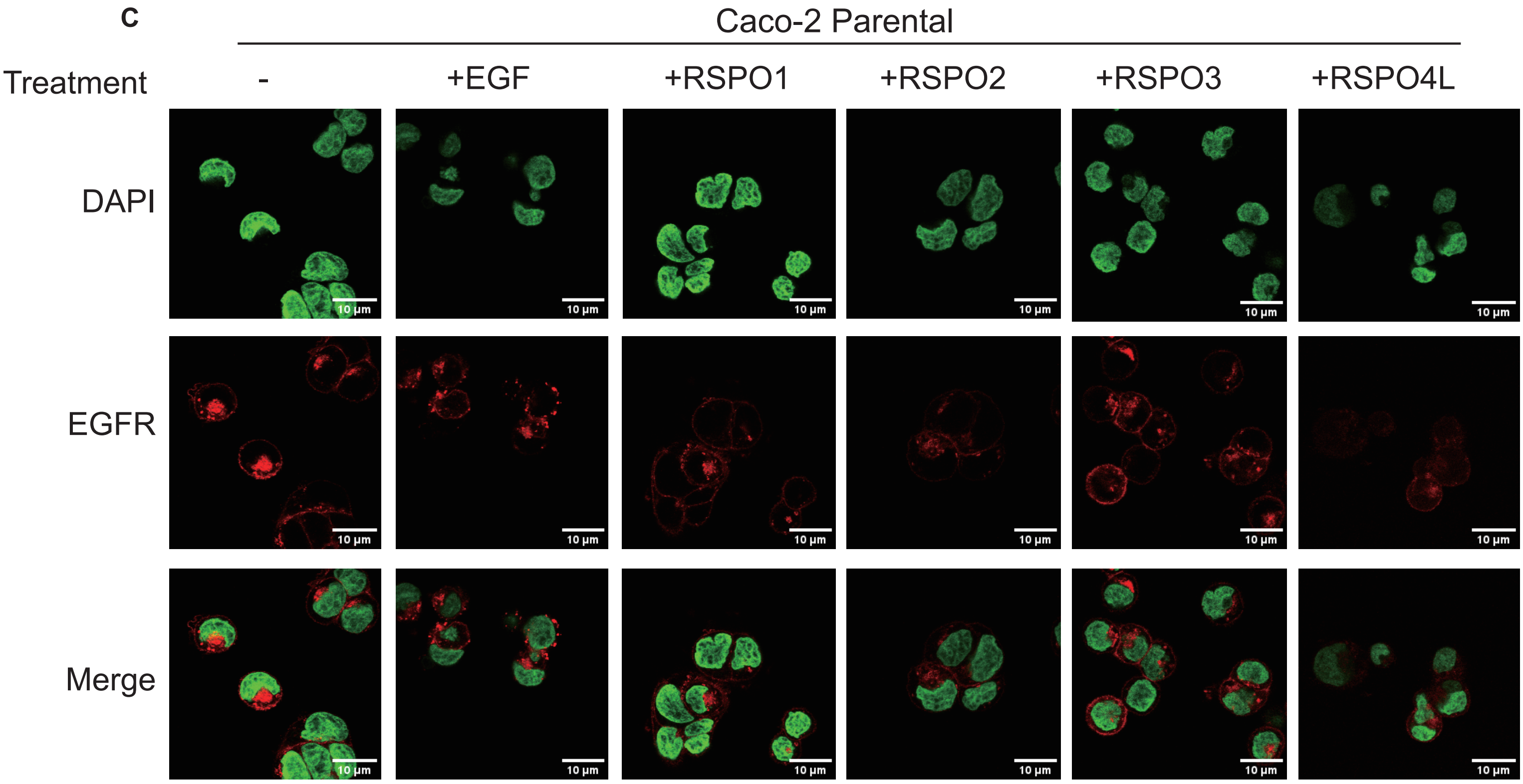


**Supplementary Fig S10.** Immunofluorescence staining of EGFR following 30 min treatment with EGF (+EGF) or RSPO-conditioned media containing RSPO1, RSPO2, RSPO3, or RSPO4L (+RSPO1/2/3/4L), as indicated. RSPO-conditioned media were collected from L cells stably expressing the corresponding RSPO variants and used at a 1:10 dilution (see Methods). Images are shown for AsPC-1 parental cells (A), AsPC-1 RNF43 KO clone A1 (B), Caco-2 parental cells (C). EGFR is shown in red and nuclei (DAPI) in green; merged images are shown in the bottom row for each condition. Scale bars, 10 µm.

Fig S11.

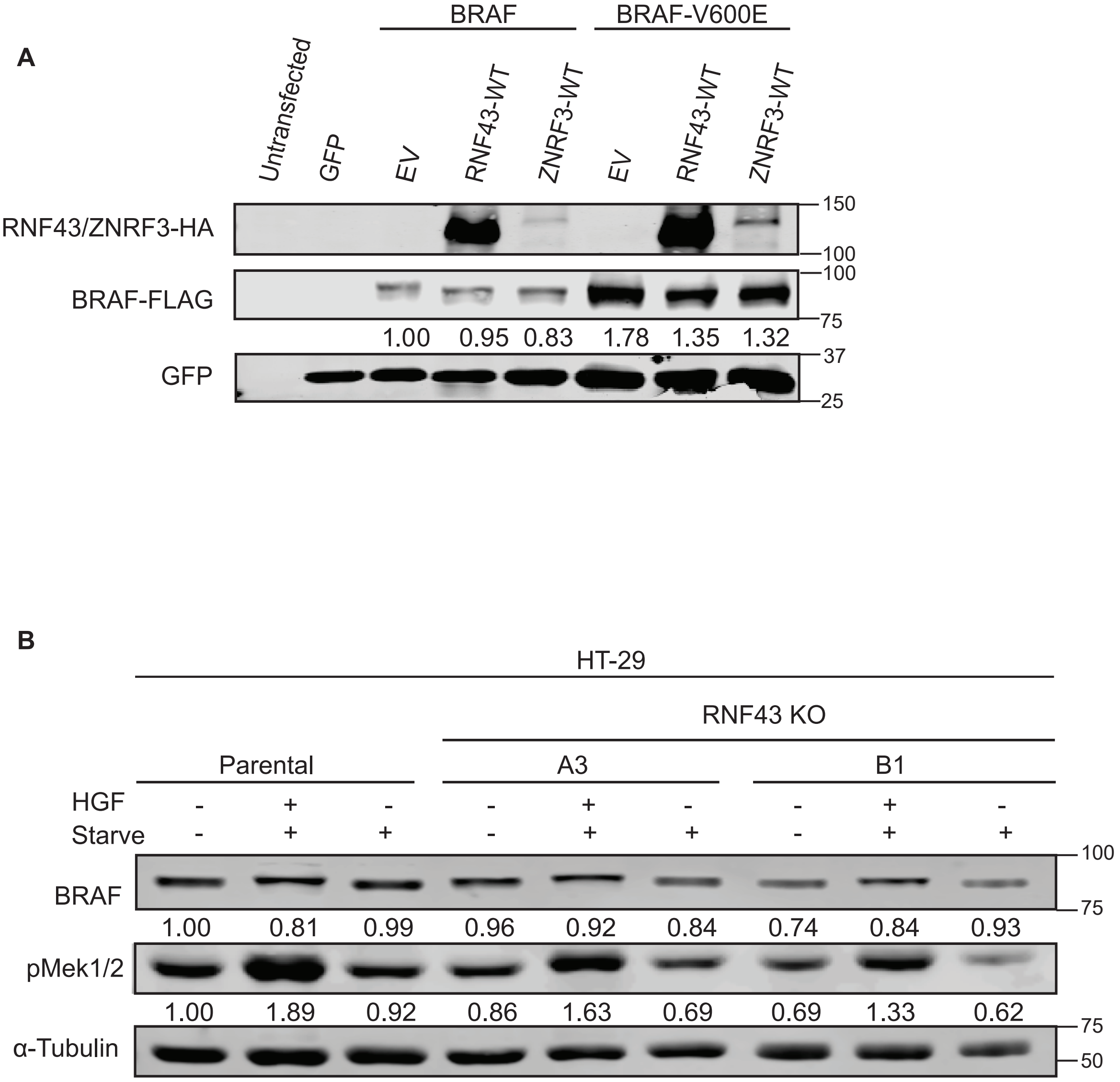


**Supplementary Fig S11.** Analysis of BRAF stability.

(A) HEK293T cells were transiently transfected with either FLAG-tagged BRAF-WT or BRAF-V600E plasmids, and co-transfected with either empty vector (EV) or plasmids expressing HA-tagged RNF43 or ZNRF3. GFP plasmid was used for normalization of transfection efficiency. Both BRAF variants are at most slightly reduced when co-expressed with RNF43 or ZNRF3.

(B) HT29 parental and RNF43-KO clones were serum-starved for 24 hours (every 3rd lane) followed by the addition of FCS-containing medium with 50 ng/ml HGF (every 2nd lane). Continuous growing HT29 cell clones were taken along as control (every 1st lane). Following starvation, re-addition of serum and HGF clearly increased pMEK1/2 levels and induced a mobility shift of BRAF indicative of multiple phosphorylation events and BRAF activation. Total BRAF levels were not affected by this treatment or RNF43 functionality status. Levels of α-Tubulin were used for normalization.

**Supplemnentary Figure S12.** Overview of Caco-2 and HT-29 cell lines carrying RNF43 knockout (KO) or wild-type (WT) alleles, with or without Cas9 integration.

| HT-29 | | | | Caco-2 | |
|---|---|---|---|---|---|
| RNF43 WT | | RNF43 KO | | RNF43 WT | |
| With Cas9 | Without Cas9 | With Cas9 | Without Cas9 | With Cas9 | Without Cas9 |
| A1 | A2 | 2B1 | A4 | A9 | E4 |
| N11 | N12 | B2 | A5 | B11 | E6 |
| 2A3 | | | | D5 | E12 |
| | | | | D8 | N4 |
| | | | | E5 | N10 |

#### HT-29 RNF43 KO Clones

Clone 2B1:

TTTGGACGCACAGGACTGGTACTGGCAGCA (Parental)
TTTGGACGGTGCTGGACGCCACCCTGATCCACCAGAGCATCACCGGCCTGTACGAGACACGGATCGACCT
GTCTCAGCTGGGAGGCGACACAGGACTGGTACTGGCAGCA (+80bp) p.T28Cfs*5
TTTGGACGAAGGGCTACAAAGAAGTGAAAAAGGACCTGATCATCAAGCTGCCTAAGTACTCCCTGTTCGA
GCTGGAAAACGGCCGgaagagaatgctggTCTGGCCCTGGCTGCTGATGGCTACCCTGCAGGCAGGCTTT
GGACGCACAGGACTGGTACTGGCAGCA (+91bp) p.T28Rfs*5

Clone B2:

CTGCAGGCAGGCTTTGGACGCACAGGACTGGTACTGGCAGCAGCGGTG (Parental)
CTGCAGG-----------------------GTACTGGCAGCAGCGGTG (-23bp) p.A23Gfs*8
CTGCAGGCAGGCTTTGGACGGGG**TAA**TTAAGACAGGAAAA... (long insertion) p.T28Gfs*2

Clone A4:

GGACGCACAGGACTGGTACTGGCA GCAGCGGTGGAGTCTGAAAGAT (Parental)
GGACG-ACAGGACTGGTACTGGCAGCAGCGGTGGAGTCTGAAAGAT (-1bp) p.T28Qfs*22

Clone A5:

CTGATGGCTACCCTGCAGGCAGGCTTTGGACGCACAGGACTGGTACTGGC (Parental)
CTGATGGC-------------------------ACAGGACTGGTACTGGC (-25bp) p.T20Qfs*22
CTGATGGCTACCCTGCAGGCAGGCTTTGGACGGCACAGGACTGGTACTGGC (+1bp) p.T28Hfs*11

### Supplementary Fig. 13

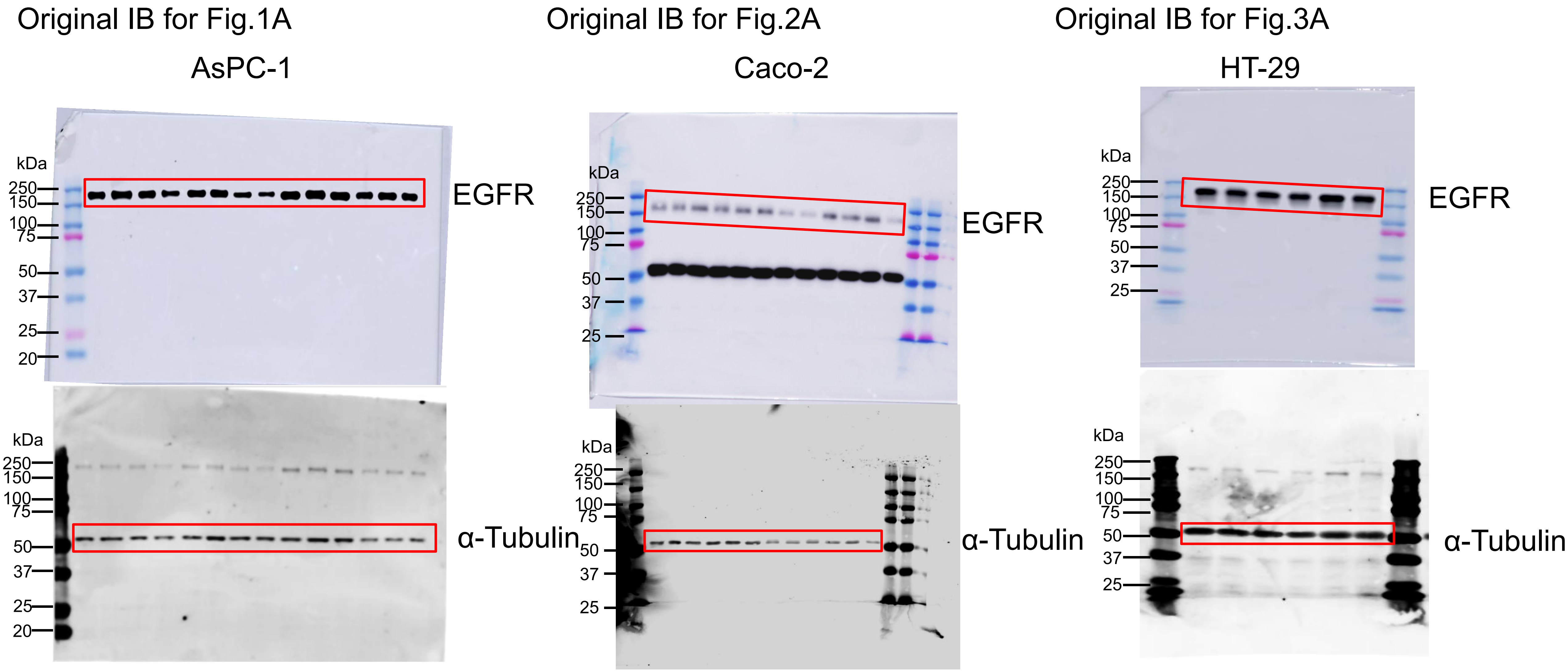

Supplementary Fig. 13, continued

Original IB for Fig.4A

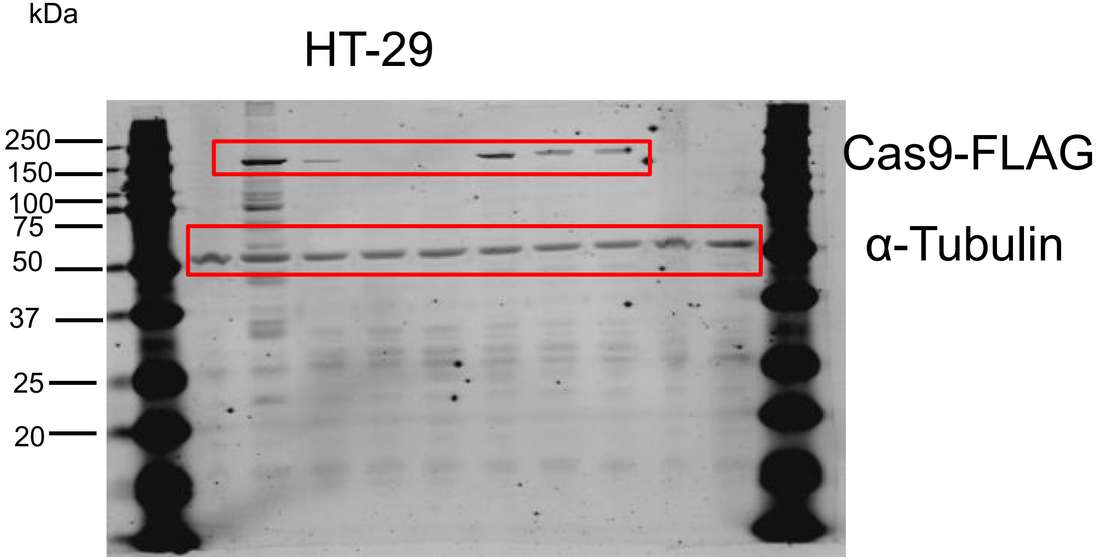


Original IB for Fig.4B

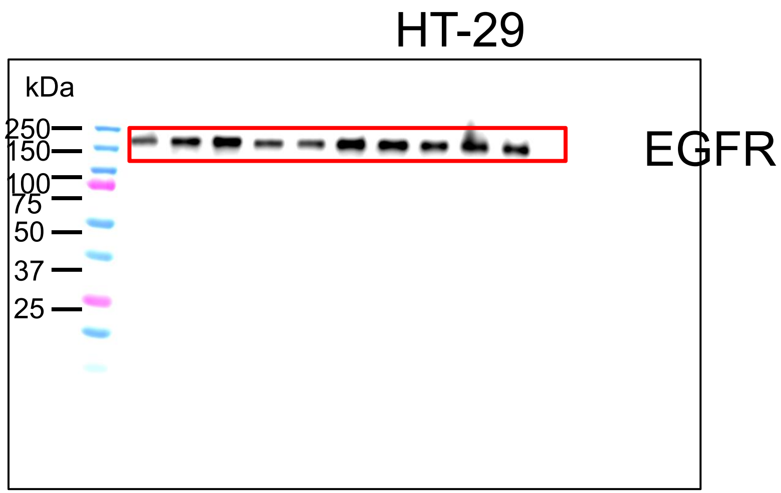


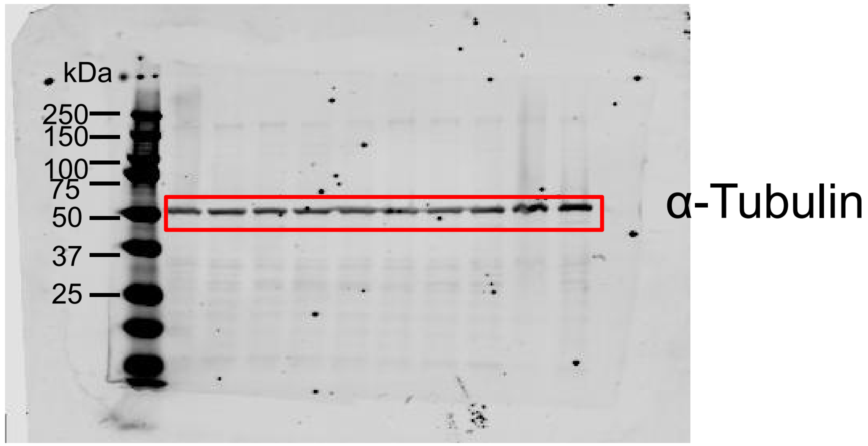

Supplementary Fig. 13, continued

Original IB for Fig.4C

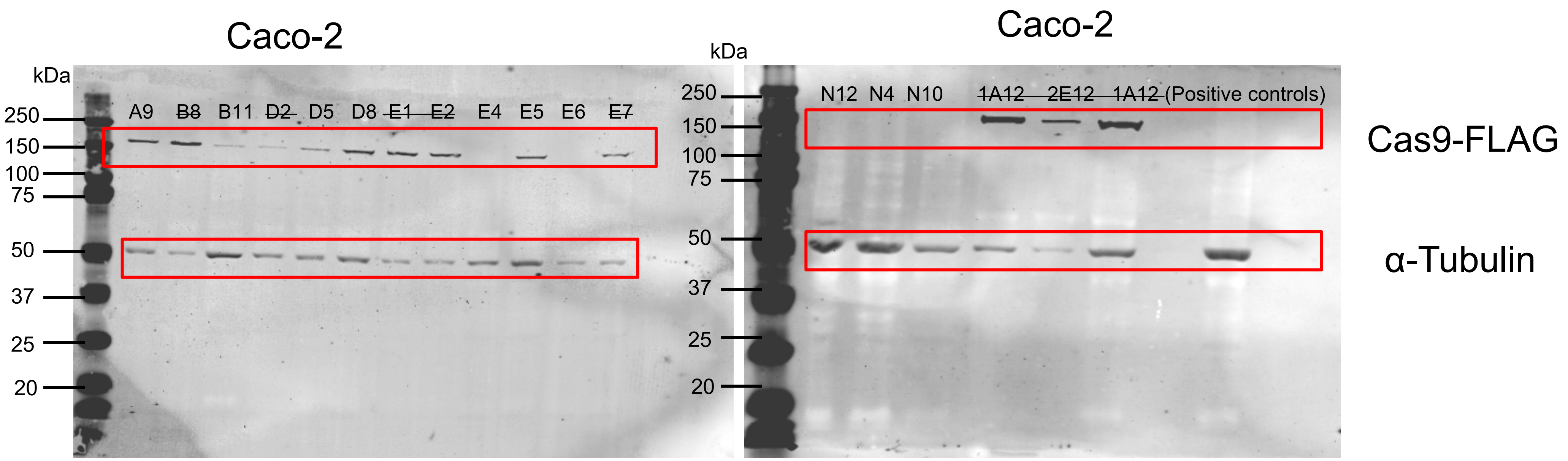

Supplementary Fig. 13, continued

Original IB for Fig.4D

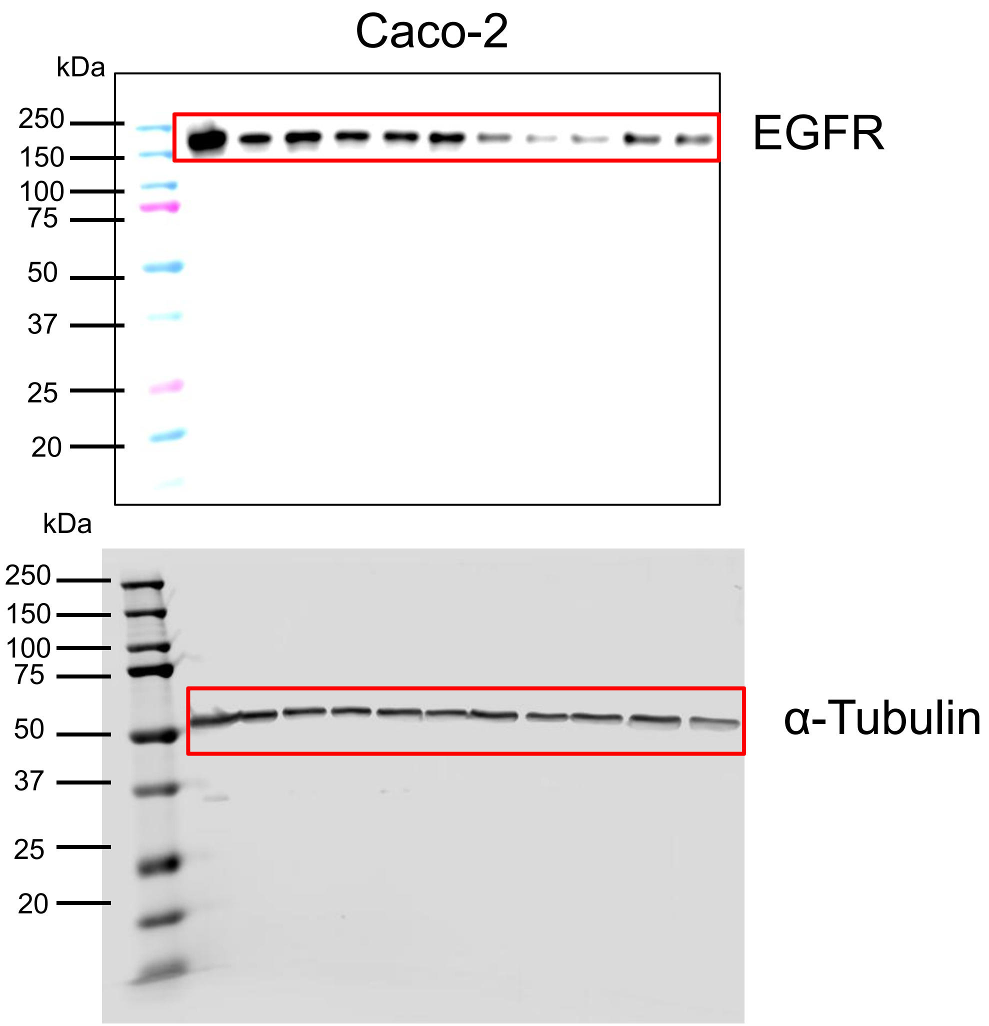

Supplementary Fig. 13, continued

Original IB for Fig.5A and 5B

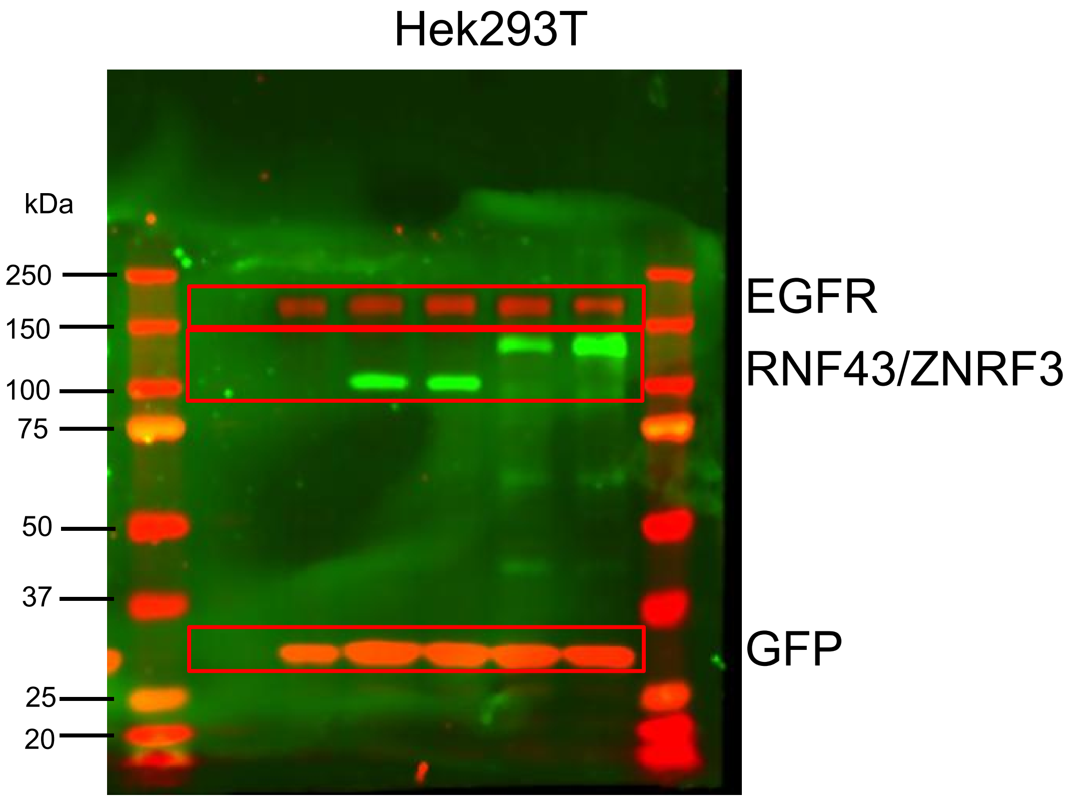


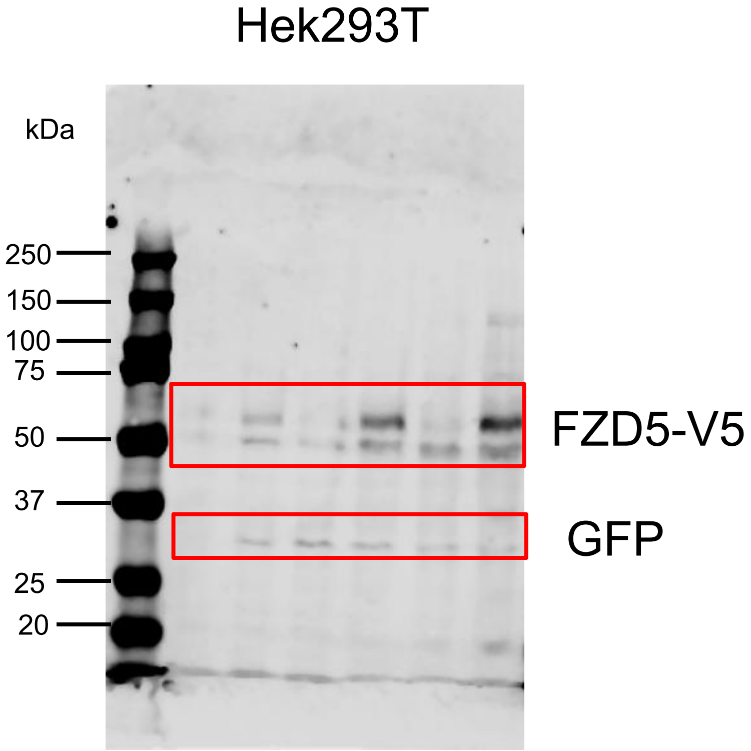

Supplementary Fig. 13, continued

Original IB for Fig.6A, 6B,6C and 6D

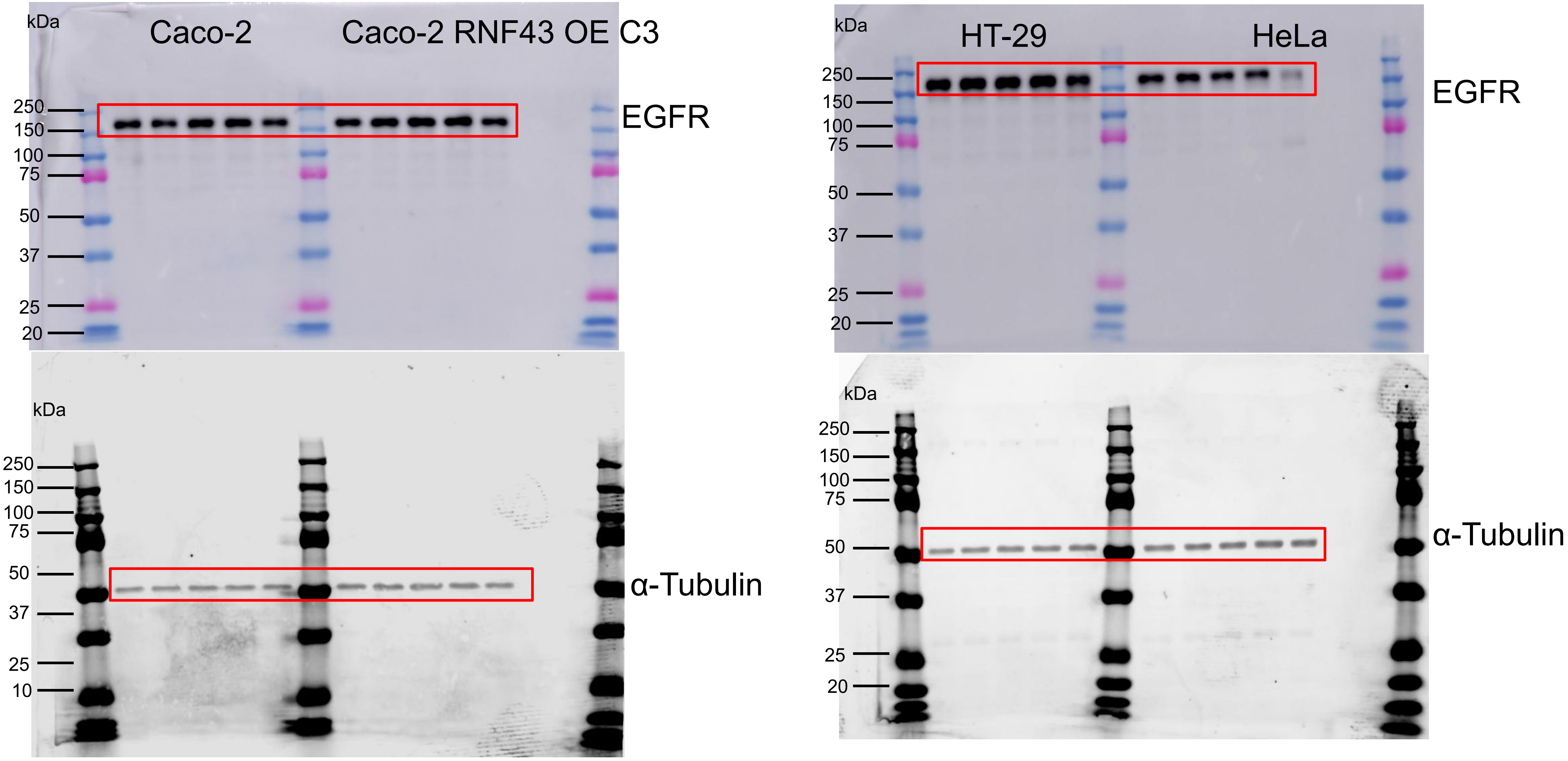

Supplementary Fig. 13, continued

Original IB for Fig. 8

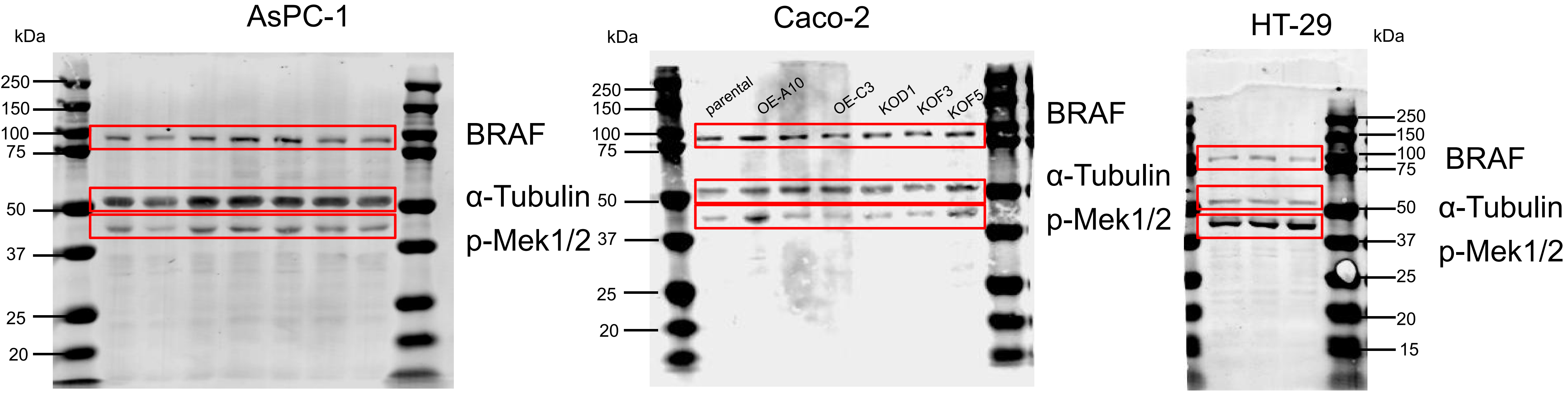

Supplementary Fig. 13, continued

Original IB for supplementary Fig.S3B
Caco-2

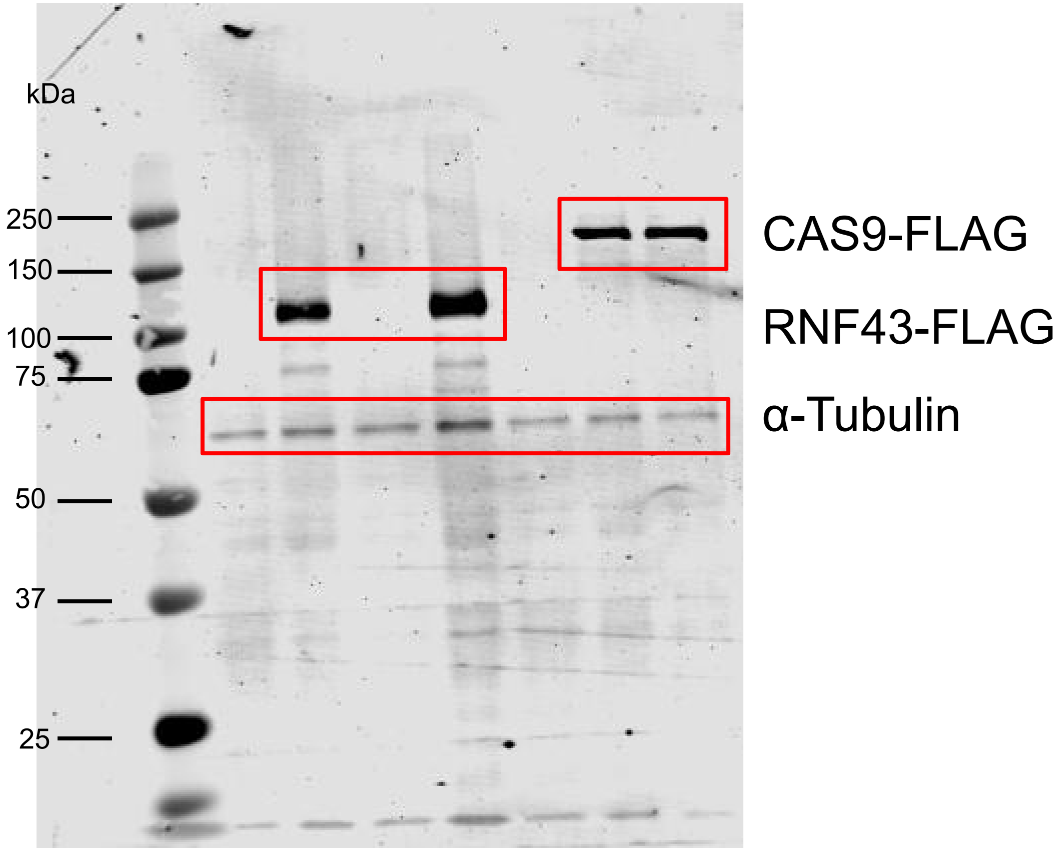


Original IB for supplementary Fig. S6

AsPC-1

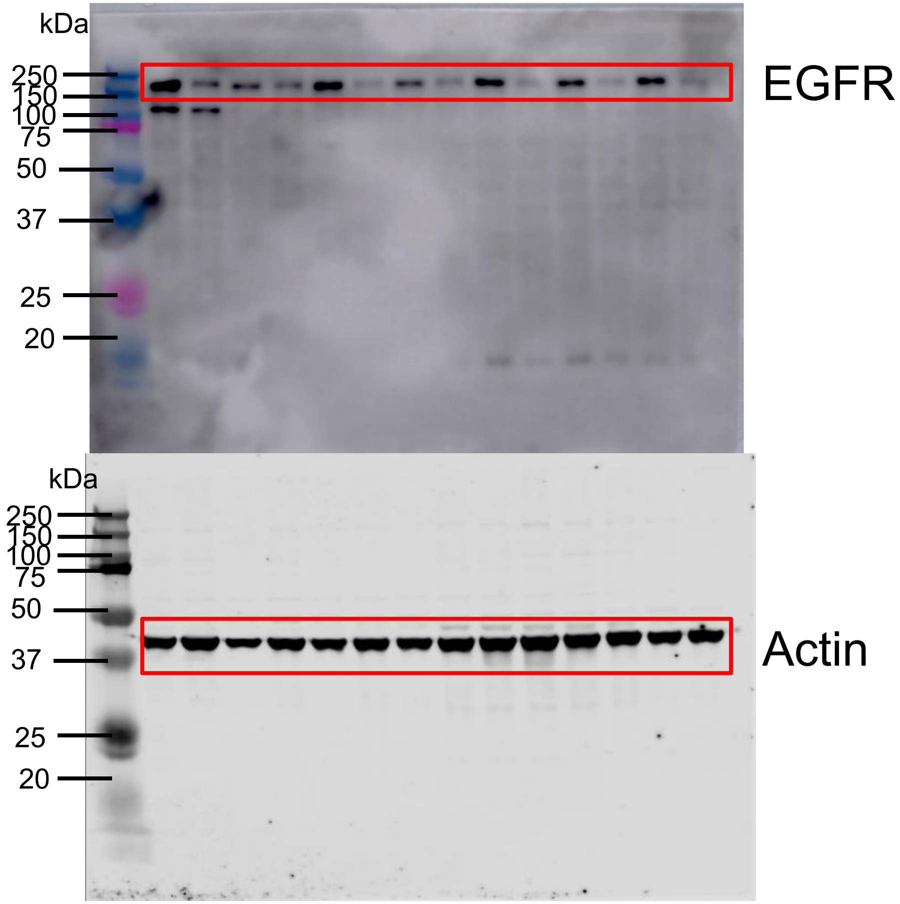

Supplementary Fig. 13, continued

Original IB for supplementary Fig. S7 A, B and C

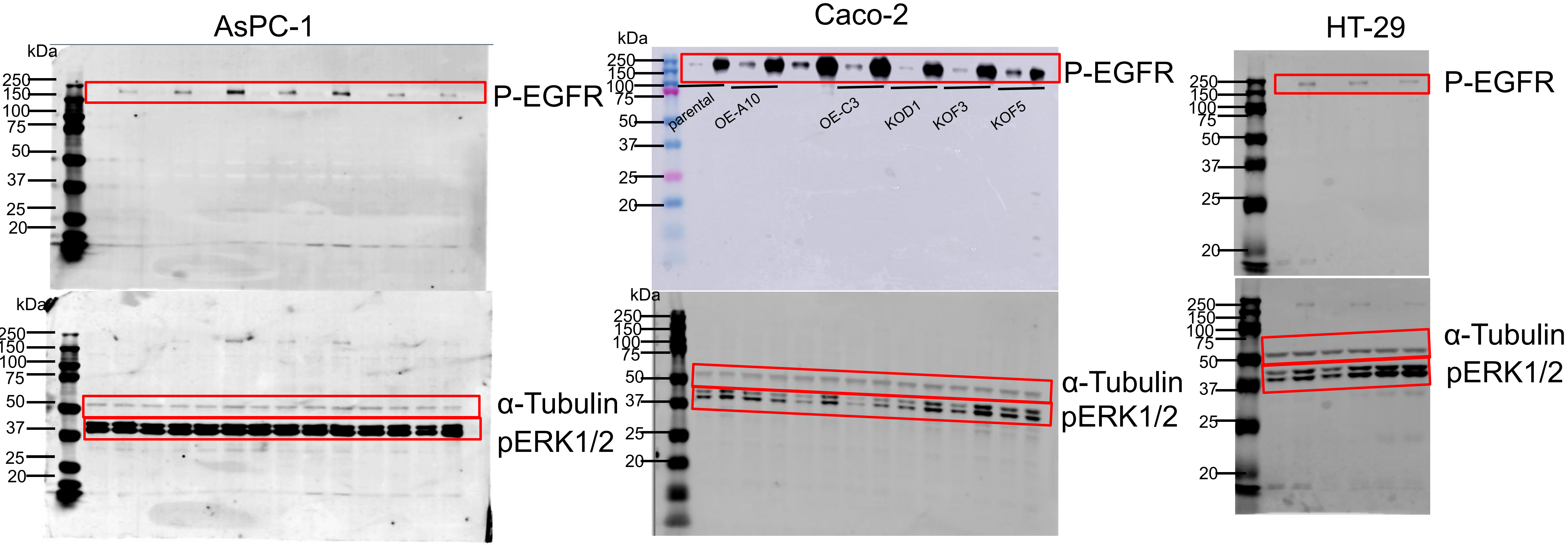

Supplementary Fig. 13, continued

Original IB for supplementary Fig.S8

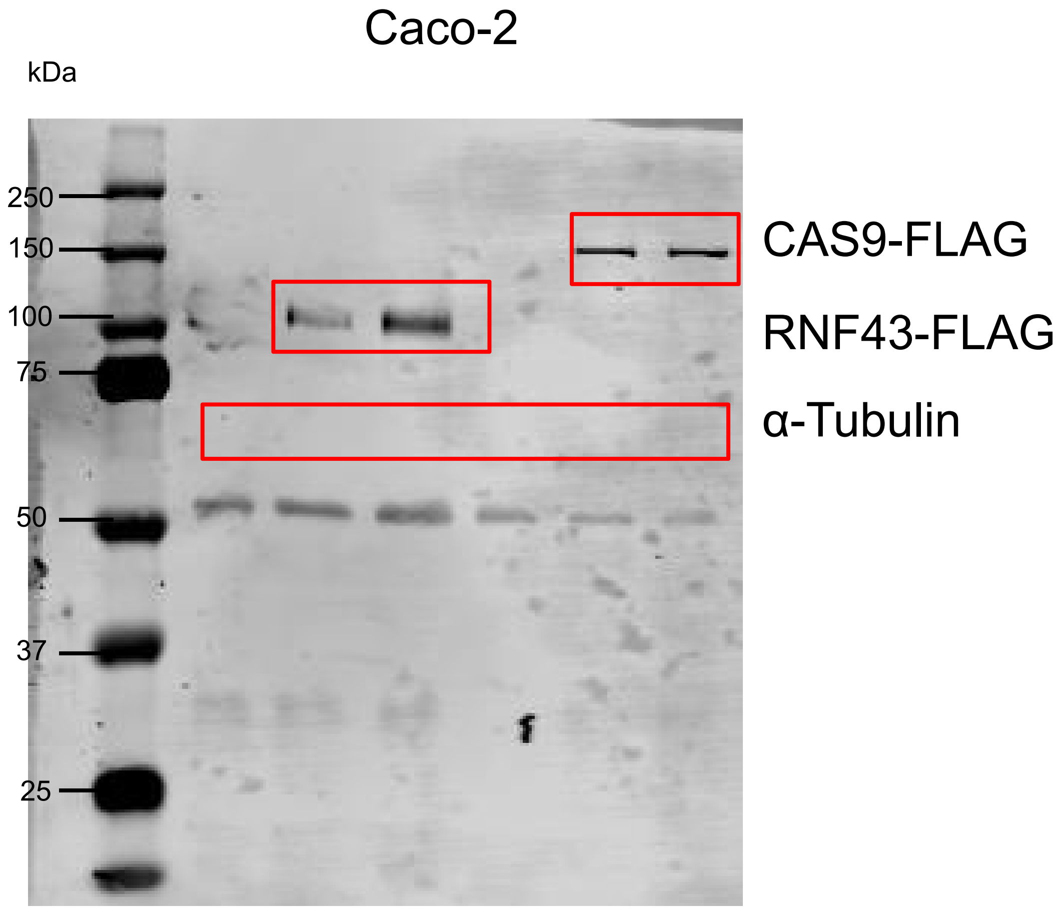

Supplementary Fig. 13, continued

Original IB for supplementary Fig.S11A

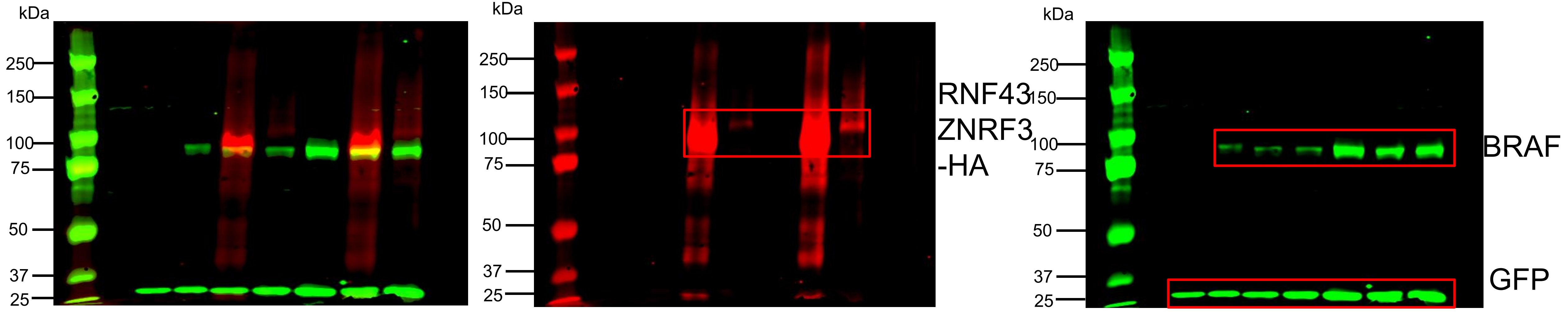


Original IB for supplementary Fig.S11B

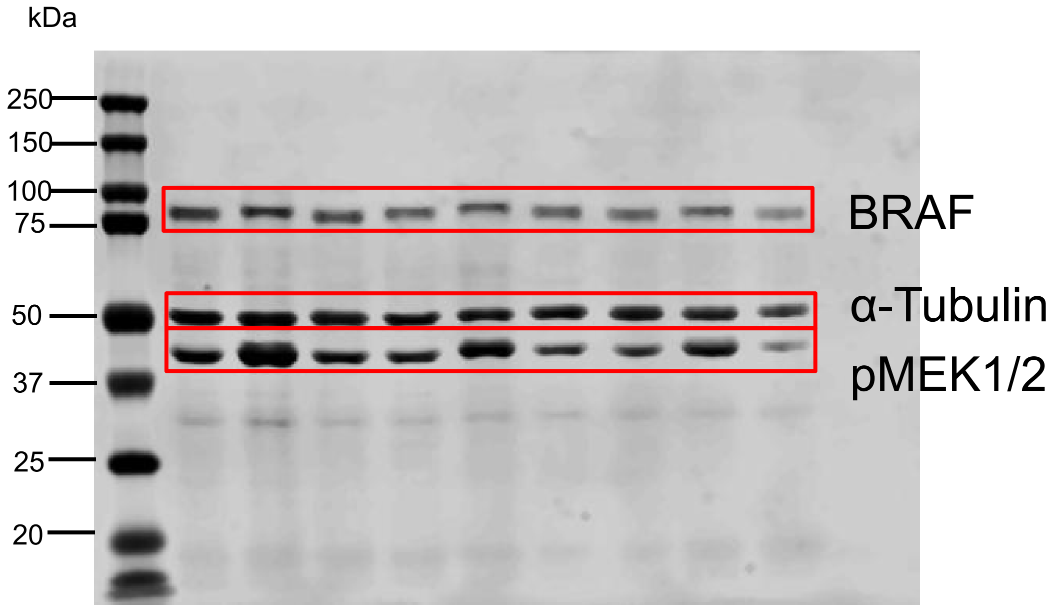

Supplementary Table S1. All Primers used in this study

| Primers | Sequences(5'-3') |
|---|---|
| Flag2HiBit-F | gttcaagaagattagcTGAGATTTCAGGTTTAAACCCG |
| Flag2HiBit-R | agccgccagccgctcacCGATCCACCTCCCGATCC |
| Rspo1_F | agccgccaccATGCGGCTTGGGCTGTGTGT |
| Rspo1_R | cacctccggaGGCAGGCCCTGCAGATGT |
| HiBiT1_F | agggcctgccTCCGGAGGTGGATCGGGAGG |
| HiBiT1_R | caagccgcatGGTGGCGGCTACAAGGCG |
| Rspo2_F | agccgccaccATGCAGTTTCGCCTTTTC |
| Rspo2_R | cacctccggaTTGGTTAGCTCTGTCTGTAG |
| HiBiT2_F | agctaaccaaTCCGGAGGTGGATCGGGAG |
| HiBiT2_R | gaaactgcatGGTGGCGGCTACAAGGCG |
| Rspo3_F | agccgccaccATGCACTTGCGACTGATTTC |
| Rspo3_R | cacctccggaGTGTACAGTGCTGACTGATAC |
| HiBiT3_F | cactgtacacTCCGGAGGTGGATCGGGAG |
| HiBiT3_R | gcaagtgcatGGTGGCGGCTACAAGGCG |
| Rspo4_F | agccgccaccATGCGGGCGCCACTCTGC |
| Rspo4_R | cacctccggaGGGCTGCAGGCCGGGCTGGC |
| HiBiT4_F | cctgcagcccTCCGGAGGTGGATCGGGAG |
| HiBiT4_R | gcgcccgcatGGTGGCGGCTACAAGGCG |
| EF1A-3XFLAG-GIB-F | acaggctgtgTCCGGAGGTGGATCGGGA |
| EF1A-3XFLAG-GIB-R | ggccaccactCATGGTGGCGGCTACAAG |
| R43-EF1A-GIB-F | cgccaccatgAGTGGTGGCCACCAGCTG |
| R43-EF1A-GIB-R | cacctccggaCACAGCCTGTTCACACAGC |
| EGFR-delGFP-F | TAAAGCGGCCGCGACTCT |
| EGFR-delGFP-R | TGCTCCAATAAATTCACTGCTTTGTG |
| Aspc-1-gRNA-3-F (HDR) | CACCGCTGTGAATTTCAGTAACAG |
| Aspc-1-gRNA-3-R (HDR) | AAACCTGTTACTGAAATTCACAGC |
| **Primers to validate BRAF plasmids** | |
| hGH-PA-R | CCAGCTTGGTTCCCAATAGA |
| BRAF-F | ACAAGGGAAAGTGGCATGGT |
| **Primers to validate RNF43 KO Clones** | |
| RNF43-ex2_cbFout | AGAGCAATGCCAAGTGATCTGA |
| RNF43-ex2_cbRout | AGCAGTAGAAGCCCGTGTAT |
| **RT-qPrimers** | **Sequences(5'-3')** |
| RNF43-ex8newF | ATCAGCATCGGACTTGCCC |
| RNF43-ex9newR | GGATGCTGGCGAATGAGGTG |

Supplementary Table S2. All antibodies used in this study

| Primary Antibobies | Species of Origin | Company | Product Number |
|---|---|---|---|
| mouse anti-HiBiT mAb(clone 30E5) | Mouse | Promega | N7200 |
| GFP Polyclonal Antibody | Rabbit | Invitrogen | A6455 |
| GFP-Tag monoclonal antibody(1G6) | Mouse | SAB | 44011 |
| HA-Tag (C29F4) | Rabbit | cell signaling technology | 3724 |
| V5-Tag (D3H8Q) | Rabbit | Cell Signaling Technology | 13202 |
| Monoclonal ANTI-FLAG M2 antibody | Mouse | Sigma | F1804 |
| Anti-alpha Tubulin antibody | Rabbit | Abcam | ab4074 |
| Anti–β-actin | Mouse | Santa Cruz Biotechnology | sc-47778 |
| EGF Receptor (D1D4J) | Rabbit | Cell Signaling Technology | 48685 |
| EGFR | Rabbit | Cell Signaling Technology | 4267 |
| EGF Receptor (D1D4J) | Rabbit | Cell Signaling Technology | 54359 |
| Phospho-EGF Receptor (Tyr1068) (D7A5) | Rabbit | Cell Signaling Technology | 3777 |
| Phospho-p44/42 MAPK (Erk1/2) (Thr202/Tyr204) | Rabbit | Cell Signaling Technology | 4370 |
| B-Raf(E3T5C) | Mouse | Cell Signaling Technology | 77622 |
| Phospho-MEK1/2 (Ser217/221) (41G9) | Rabbit | Cell Signaling Technology | 9154 |
| **Secondary Antibodies** | **Species of Origin** | **Company** | **Product Number** |
| IRDye® 800CW Goat anti-Rabbit IgG Secondary Antibody | Goat | LI-COR | 926-32211 |
| IRDye® 680CW Goat anti-Rabbit IgG Secondary Antibody | Goat | LI-COR | 926-68071 |
| IRDye® 800CW Goat anti-Mouse IgG Secondary Antibody | Goat | LI-COR | 926-32210 |
| IRDye® 680CW Goat anti-Mouse IgG Secondary Antibody | Goat | LI-COR | 926-68070 |
| Anti-rabbit IgG, HRP-linked Antibody | Goat | Cell Signaling Technology | 7074 |
